# Impaired p53-mediated DNA damage response contributes to microcephaly in Nijmegen Breakage Syndrome patient-derived cerebral organoids

**DOI:** 10.1101/2020.09.29.318527

**Authors:** Soraia Martins, Lars Erichsen, Angeliki Datsi, Wasco Wruck, Wolfgang Goering, Krystyna Chrzanowska, James Adjaye

**Affiliations:** Institute for Stem Cell Research and Regenerative Medicine, Medical Faculty, Heinrich-Heine University, 40225 Düsseldorf, Germany; Institute for Transplantation Diagnostics and Cell Therapeutics, Heinrich-Heine University, 40225 Düsseldorf, Germany; Institute for Pathology, University Hospital and Medical Faculty of the Heinrich-Heine University Düsseldorf, 40225 Düsseldorf, Germany; Department of Medical Genetics, Children’s Memorial Health Institute, Warsaw, Poland

**Keywords:** NBS, iPSCs, cerebral organoids, disease modelling

## Abstract

Nijmegen Breakage Syndrome (NBS) is a rare autosomal recessive genetic disorder caused by mutations within *NBN*, a DNA-damage repair protein. Hallmarks of NBS include several clinical manifestations such growth retardation, chromosomal instability, immunodeficiency and progressive microcephaly. However, the etiology of microcephaly in NBS patients remains elusive. Here, we employed induced pluripotent stem cell-derived brain organoids from two NBS patients to analyze the underlying mechanisms of microcephaly. We show that NBS-organoids carrying the homozygous 647del5 *NBN* mutation are significantly smaller in size with disrupted cyto-architecture Patient-derived organoids exhibit premature differentiation together with neuronatin (NNAT) overexpression and key pathways related to DNA damage response and cell cycle are differentially regulated compared to controls. Moreover, we show that after exposure to bleomycin, NBS organoids undergo a delayed p53-mediated DNA damage response and aberrant trans-synaptic signalling, which ultimately leads to neuronal apoptosis. Our data provide insights into how mutations within *NBN* alters neurogenesis in NBS patients, thus providing a proof of concept that cerebral organoids are a valuable tool for studying DNA damage-related disorders.

## Introduction

Development of the nervous system is a strictly regulated process whereby neural progenitor cells (NPCs) rapidly proliferate and potentially generate high-levels of oxidative stress. Neurons albeit being post-mitotic cells display high rates of metabolism and mitochondria activity also contributing to a stressful environment which increases the susceptibility to DNA damage. Deficiency in the DNA damage response (DDR) causes many syndromes with pronounced neuropathology [1,2]. Hypomorphic mutations within *nibrin* (*NBN*), coding a DDR protein, leads to Nijmegen Breakage Syndrome (NBS), a rare autosomal recessive genetic disorder belonging to the chromosomal instability syndromes [3]. NBS was first described in 1979 at the University of Nijmegen and it has been considered as a multisystemic disorder [4]. Clinically, NBS is characterized by severe and progressive microcephaly, growth retardation, typical facial appearance, premature ovarian failure, deterioration of the cognitive functions, immunodeficiency, chromosomal instability and elevated sensitivity to ionizing radiation. Apart from microcephaly, other developmental abnormalities of the brain have been reported in a few patients, including neuronal migration disorder, agenesis of the corpus callosum and arachnoid cysts [5–7]. By the age of 20 years, more than 40% of NBS patients develop malignancy diseases, predominantly of hematological origin which, in addition to the recurrent infections are the major cause of death in these patients [8].

The worldwide prevalence is estimated at 1:100000 live births, however NBS is particularly common in Eastern Europe with carrier frequencies as high as 1:155 in some populations of Czech Republic, Poland and Ukraine. [9]. More than 90% of the NBS patients are homozygous for a founder mutation, a five base pair deletion in exon 6 (657del5) within *NBN*. Due to alternative translation from a cryptic start site upstream of the deletion, this mutation leads to the truncation of the wild-type protein into two different fragments: a 26 kDa amino-terminal protein (p26) and a 70 kDa carboxy-terminal (p70), which retain some residual functions [10]. NBN together with MRE11 and RAD50 form the MNR complex which plays a central role in DNA damage signaling and repair, telomere maintenance, proper centromere duplication, cell cycle checkpoint activation and processing of stalled replication forks [11].

With NBN playing a multifunctional central role, it is not surprising that lymphoblasts and fibroblasts from NBS patients exhibit chromosomal instability with impaired cell cycle and regulation of apoptosis [12–14]. However, the functional consequences of *NBN* 657del5 during neurodevelopment remained largely unexplored. To this end several studies based on *Nbn* conditional knockdowns have been performed [15–20]. While these studies provided helpful information, some of the mouse models do not reproduce the brain-related phenotype seen in patients and in the ones that the microcephaly phenotype was present, divergent results were found. Frappart *et. al* (2005) showed inactivation of *Nbn* in the mouse neuronal tissue resulted in decreased proliferation of the NPCs and increase apoptosis of postmitotic neurons in the cerebellum, by activation pf p53 [20]. However, Zhou *et al.* (2012) showed that deletion of *Nbn* in the mouse central nervous system affects proliferation and apoptosis mainly in the cortex ventricular zone proliferating area (VZ), affecting only NPCs [19].

To bridge the gap between the mouse models and the human phenotype of NBS, induced pluripotent stem cells (iPSCs) technology provides the best platform to derive a reliable human disease model to study the effects of *NBN* mutations in neurons derived from NBS patients. In fact we previously reported that fibroblasts from NBS patients can be reprogrammed into iPSCs and thus by-passing premature senescence [21]. Global transcriptome analysis of NBS fibroblasts and NBS-iPSCs unveiled de-regulated cancer related pathways such as p53, cell cycle and glycolysis [22]. Differentiation of the NBS-iPSCs into NPCs showed these cells have de-regulated expression of neural developmental genes in-part due to NBS-NPCs inability to maintain normal levels of p53 [23]. Recent advances in *in vitro* culture of 3D cerebral organoids derived from iPSCs have illuminated the early mechanisms of mammalian neurodevelopment. Cerebral organoids provide a unique opportunity to model human organogenesis through the presence of an organizational feature unique to 3D brains, such as cortical layers [24–26]. Indeed, studies have shown the utility of cerebral organoids from patients-derived iPSCs to unveil the mechanisms of microcephaly and other neurodevelopmental disorders [27].

Here, we took advantage of the cerebral organoids system to analyze the underlying mechanisms of microcephaly and other brain abnormalities present in NBS patients. NBS-organoids carrying the homozygous 647del5 *NBN* mutation are significantly small with disrupted cyto-architecture. Both patient-derived organoids exhibit premature differentiation and key pathways related to DNA damage response and cell cycle are differentially regulated compared to controls. After exposure to DNA damage, NBS organoids present a delay in the activation of cell cycle arrest due to inability to stabilize p53. Our data provide insights into how hypomorphic mutations within *NBN* alters neurogenesis in the NBS patients, thus providing a proof of concept that cerebral organoids are a valuable tool enabling research into DNA damage-related disorders.

## Results

### iPSCs-derived from NBS patients exhibit chromosome instability

In order to create an *in vitro* model system to study NBS, we previously generated iPSC lines from two NBS patients, referred here as NBS1 and NBS8-iPSCs [21,28]. NBS1-iPSCs carry the homozygous 657del5 mutation and NBS8-iPSCs the heterozygous 657del5 mutation. As NBS is characterized by chromosomal instability, genome integrity evaluation was performed at the beginning of the current study. While NBS1-iPSCs presents a normal karyotype (**Table S1, Figure S1B**), copy number variations (CNV) analysis using array comparative genome hybridization (array-GCH) showed that, although no larger chromosomal aberrations was reported, there is a loss of heterozygosity (LOH) in chromosome 8q, region where *NBN* is located **(Supplementary figure 1A)**. Of note, the detected LOH was pre-existing in the parental fibroblast line. Karyotype analysis of NBS8-iPSCs revealed a higher number of acquired CNVs and cytogenetic rearrangements, namely a duplication of most of the long arm of chromosome 5 and a duplication of the telomeric end of chromosome 3p **(Figure S1B-C)**. These aberrations were acquired during the reprogramming process, as NBS8-iPSC line was generated by retroviral-based reprogramming.

**Table S1** presents the detailed information about the iPSC lines used in this study. As controls, HFF-derived iPSCs [29], here referred as CTR1-iPSCs and urine cell-derived iPSCs [30], here referred as CTR2-iPSCs, were used.

### NBS-iPSCs efficiently differentiate into cerebral organoids and recapitulate the microcephaly phenotype

To address how human early neurodevelopment is affected in NBS and to dissect the mechanisms underlying microcephaly in NBS patients, we implemented a publish suspension protocol to generate forebrain organoids. An equal number of dissociated iPSCs were aggregated in a minimally pattern conditions and in the absence of extracellular scaffolding [31]. To facilitate neuralization, iPSCs were cultured in the presence of two inhibitors of the SMAD pathway, followed by a proliferation and corticogenesis progression phase due to the addition of EGF and FGF. The final maturation phase was achieved by exposing the cerebral organoids to BDNF. In order to increase oxygen and nutrient diffusion, cerebral organoids were kept under orbital shaking conditions **(Figure 1A)**.

By day 20, immunohistochemical analysis revealed that control- and NBS-organoids showed an internal cyto-architecture composed mostly of the NPCs expressing SOX2 aligned in a VZ-like structure surrounded by the DCX+ neuroblasts that will gradually generate the neurons from the cortical plate **(Figure 1B)**.

**Figure 1.**
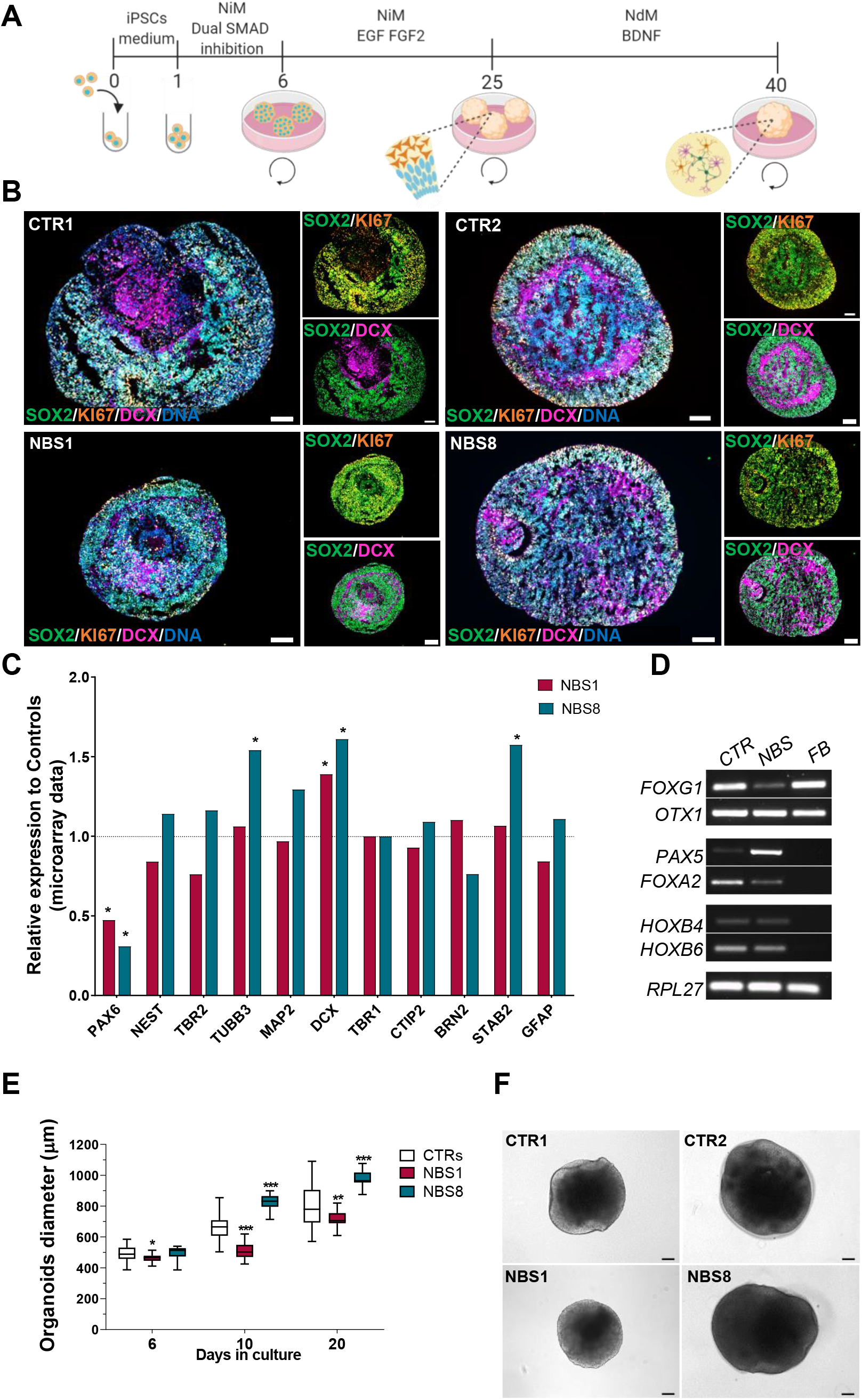
Generation and characterization of NBS cerebral organoids. **(A)** Schematic outline of the main stages of the differentiation protocol to generate the iPSC-derived cerebral organoids. **(B)** Representative immunocytochemistry images of the distribution of cells expressing SOX2, KI67 and DCX in cerebral organoids at day 20. Scale bars, 100μm. **(C)** Relative mRNA expression analysis of progenitor markers (*PAX6*, *NES* and *TBR2*), pan-neuronal makers (*TUBB3*, *MAP2*, *DCX*), early-born neurons (*TRB1*, *CTIP2*), late-born neurons (*BRN2* and *STAB2*) and the astrocytes marker *GFAP* in NBS-organoids (NBS1 and NBS8) compared to control-organoids (CTR1 and CTR2). **(D)** RT-PCR analysis for brain-region specificity at day 20 in control- and NBS-organoids (forebrain: *FOXG1* and *OTX1*; midbrain: *PAX5* and *FOXA2* and hindbrain: *HOXB4* and *HOXB6*). FB, fetal brain control. **(E)** Comparison of the diameter of control- (n=76 day 6, n=61 day 10 and n=55 day 20 for both CTR1 and CTR2), NBS1- (n=15 day 6, n=22 day 10 and n=18 day 20) and NBS8-organoids (n=36 day 6, n=35 day 10 and n=33 day 20) cerebral organoids at day 6, 10 and 20. The diameter was significantly smaller in NBS1-organoids. Significance in comparison to control (CTR1 and CTR2) was calculated with one-way ANOVA followed by Dunnett’s multiple comparison test; *p<0.05, **p<0.01, ***p<0.001. **(F)** Representative bright-field images of control and NBS-cerebral organoids at day 20. NBS1 organoids visually lack neuroepithelial structures. Scale bars, 100μm.

To evaluate the efficiency of the differentiation, mRNA levels of progenitor markers, pan-neuronal makers, early born neurons, late-born neurons and the astrocytes marker *GFAP* were assessed. While there is some heterogeneity between the expression of the neuronal markers between NBS1 and NBS8 organoids, neuronal progenitor markers such as *PAX6* were significantly down-regulated in NBS organoids. On the other hand, NBS organoids showed an up-regulation of the neuronal markers *DCX* and *TUBB3* **(Figure 1C)**.

To test the regional specificity of the cerebral organoids, we performed a RT-qPCR for forebrain, midbrain and hindbrain markers. Despite the use of a protocol that was developed to yield region specific forebrain organoids, both 20 days control and NBS cerebral organoids expressed not only the forebrain markers *FOXG1* and *OTX1*, but also the midbrain markers *PAX5* and *FOXA2* and the hindbrain markers *HOXB4* and *HOXB6,* albeit at lower expression levels **(Figure 1D)**.

To analyse the developmental stage of the cerebral organoids with respect to human fetal brain, we compared the transcriptome profile of CTR1, CTR2, NBS1 and NBS8-organoids at day 20 to the transcriptomic data from the Allen Human Brain Atlas (https://www.brainspan.org) [32]. Our cerebral organoids at day 20 were closely related to 8-9 post-conception week (pcw) **(Figure S2C)**. To test if NBS-organoids could recapitulate the microcephaly phenotype we evaluated the size of the organoids over a period of 20 days. Although the size of NBS8-organoids was similar to that of controls at day 6, during prolonged culture the organoids became significantly larger. On the other hand, NBS1-organoids were significantly smaller in comparison to controls and the size was not compensated during the time in culture **(Figure 1E)**. In addition to the smaller size, NBS1-organoids were relatively spherical and visually lacked the neuroepithelial structures observed in the controls and NBS8-organoids **(Figure 1F)**.

Together these results imply that control- and NBS-iPSCs efficiently differentiate into cerebral organoids, which recapitulate aspects of early human brain development with defined molecular markers. Furthermore, NBS1-organoids, carrying the homozygous *NBN* mutation, recapitulate the microcephaly phenotype.

### NBS organoids present a disrupted cyto-architecture with normal proliferation of the NPCs

NPCs display incredible plastic features and an enhanced capacity to sense the environment. NPCs can divide both symmetrically and asymmetrically, stay quiescent for long periods, can undergo active proliferation or differentiate upon instruction. Environmental cues can trigger the proliferation of NPCs or the acquisition of a terminally differentiated phenotype [33]. We analysed the expression of the NPCs marker SOX2 together with the proliferation marker *KI67* in our cerebral organoids **(Figure 2A)**. At day 20 both control- and NBS-organoids were mainly composed of SOX2^+^ NPCs, with no significant differences in the total number of SOX2^+^ cells **(Figure 2B)** as well as in the *SOX2* mRNA expression **(Figure 2C)** between control- and NBS-organoids. Furthermore, a similar proliferation of the NPCs was observed based on KI67 expression **(Figure 2D-F)**.

**Figure 2.**
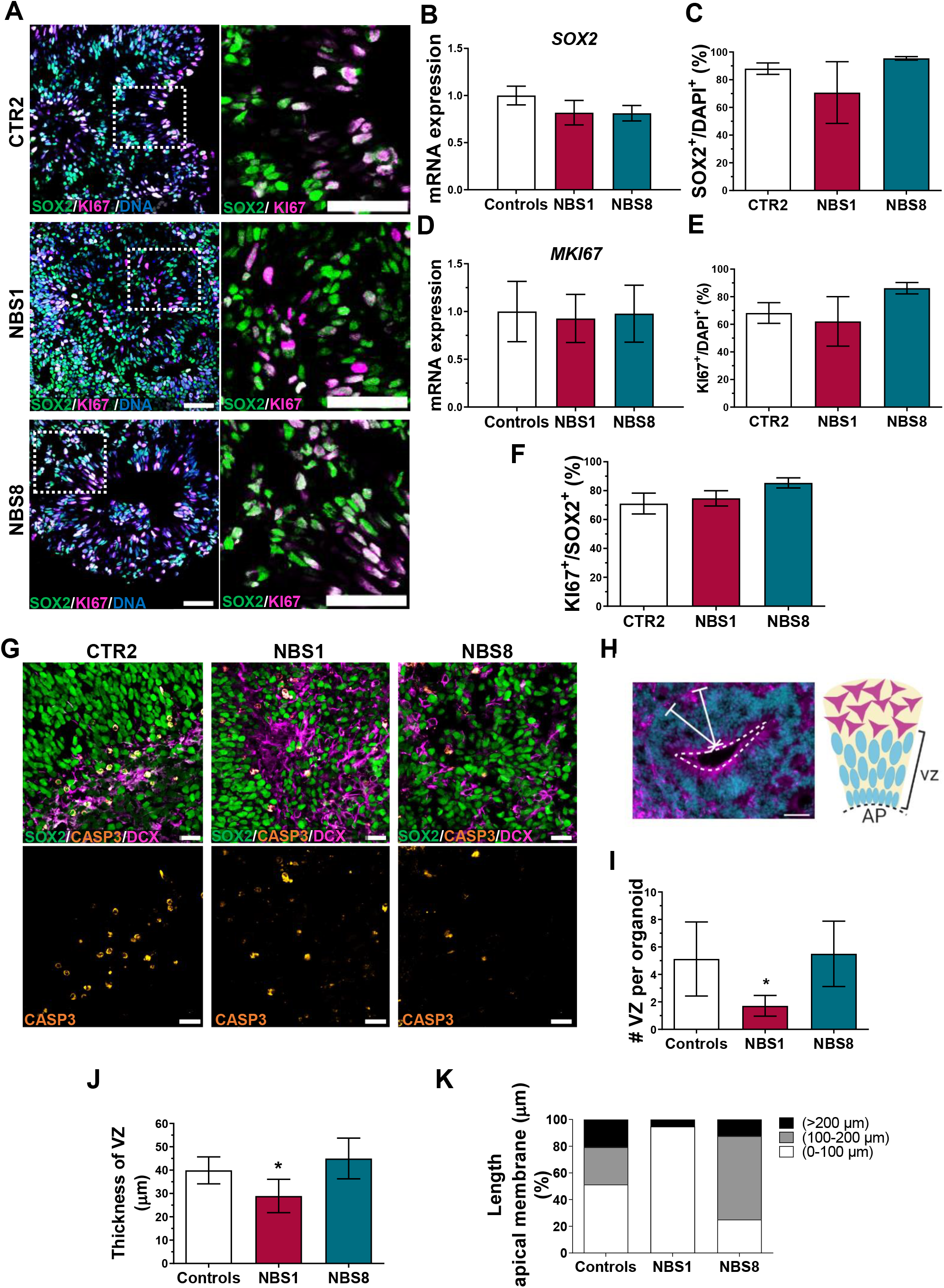
Analysis of proliferation of the NPCs and VZs cyto-architecture at day 20. **(A)** Representative confocal pictures of immunostainings for SOX2 and KI67 in CTR2-, NBS1- and NBS8-organoids. Scale bars, 50μm. **(B)** qRT-PCR analysis of *SOX2* mRNA expression in NBS1 and NBS8-organoids relative to control organoids (CTR1 and CTR2). Results are mean ± 95% confidence interval derived from 3 independent differentiations. **(C)** Quantification of the SOX2-positive cells in CTR2 and NBS1- and NBS8-organoids. Results are mean ± SD derived from 3 organoids from independent differentiations. **(D)** qRT-PCR analysis of *MKI67* mRNA expression in NBS1 and NBS8-organoids relative to control organoids (CTR1 and CTR2). Results are mean ± 95% confidence interval derived from 3 independent differentiations. **(E)** Quantification of the KI67-positive cells in CTR2 and NBS1- and NBS8-organoids. Results are mean ± SD derived from 3 organoids from independent differentiations. **(F)** Quantification of the KI67-positive cells within the SOX2-positive cells in CTR2-, NBS1- and NBS8-organoids. Results are mean ± SD derived from 3 organoids from independent differentiations. **(G)** Representative confocal pictures of immunostainings for SOX2, cleaved-CASP3 and DCX in CTR2-, NBS1- and NBS8-organoids. Cleaved-CASP3 colocalized with the DCX positive cells. Scale bars, 50μm. **(H)** Schematic illustration of a ventricular zone (VZ) and how the thickness of the VZ and the length of the apical membrane were calculated in order to evaluate the cyto-architecture of the cerebral organoids. **(I)** Quantification of the number of the VZs per organoid in control (CTR1 and CTR2) and NBS1 and NBS8-organoids. Results are mean ± SD derived from 3 organoids from independent differentiations. Significance in comparison to control was calculated with one-way ANOVA followed by Dunnett’s multiple comparison test. *p<0.05. **(J)** Quantification of the thickness of the VZs in μm in control (CTR1 and CTR2) and NBS1 and NBS8-organoids. Results are Results are mean ± SD derived from 3 organoids from independent differentiations. Significance in comparison to control was calculated with one-way ANOVA followed by Dunnett’s multiple comparison test. *p<0.05 **(K)** Quantification of the length of the apical membrane per VZ in control-, NBS1- and NBS8-organoids; 3 organoids from independent differentiations were analysed.

Next, we examined if the reduction in size observed in NBS1-organoids was due to enhanced cell apoptosis in the cerebral organoids. We found a comparable number of cleaved Caspase-3^+^ cells between control and NBS organoids, mostly located outside the VZ and co-localized with the DCX^+^ cells. Lastly, we explored the internal cyto-architecture of the cerebral organoids by determined the number and the thickness of the VZ, as well as the length of the apical membrane. While control organoids displayed typical VZ containing SOX2^+^ NPCs proliferating at the apical surface of the ventricular zone **(Figure 2H)**, NBS1-organoids exhibited a markedly reduction of these well-defined VZs (Figure 2I). Likewise, the VZs present were significant smaller (Figure 2J) with smaller apical membrane **(Figure 2K)**. Notably, at day 20 there were no clear differences in the VZs from the NBS8-organoids compared to control.

Our results suggest that although the population of the NPCs is not affected in an early developmental stage, NBS1-organoids can show a disrupted cyto-architecture that can affect normal brain development.

### NBS1 and NBS8 organoids show a distinct transcriptomic profile

To gain further insights into the pathophysiology of NBS, we performed transcriptome analysis of both control and NBS organoids at day 20 of differentiation. We then applied bioinformatic analysis to our transcriptome data to perform a cluster analysis and to identify the differentially expressed genes (DEGs). Hierarchical clustering revealed one cluster containing the NBS8-organoids and the second cluster containing CTR1-, CTR2- and NBS1-organoids, thus indicating that NBS8-organoids have a distinct transcriptome profile **(Figure S2A-B)**.

To further identify the DEGs, we compared the transcriptome of the NBS1- and NBS8-organoids to both CTR-organoids. Among the commonly expressed genes between NBS1 and controls, we found 198 up-regulated and 210 down-regulated DEGs **(Figure 3A, Table S2)**. Enrichment analysis of the up-regulated DEGs showed *subpallium development*, *regulation of focal adhesion assembly*, *phospholipid dephosphorylation* and *centromere complex assembly* as the most enriched GOs **(Figure 3B)**. Regarding the canonical pathways, up-regulated DEGs were significantly enriched in *RTMs methylate histone arginines*, *Signalling by GPCR*, *DNA damage/Telomere stress induce senescence* and *Cellular senescence* **(Figure 3C)**. Down-regulated genes were enriched for GOs as *chemotaxis*, *negative regulation of proteolysis* and *cellular response to hormonal stimuli* **(Figure 3D)**.

**Figure 3.**
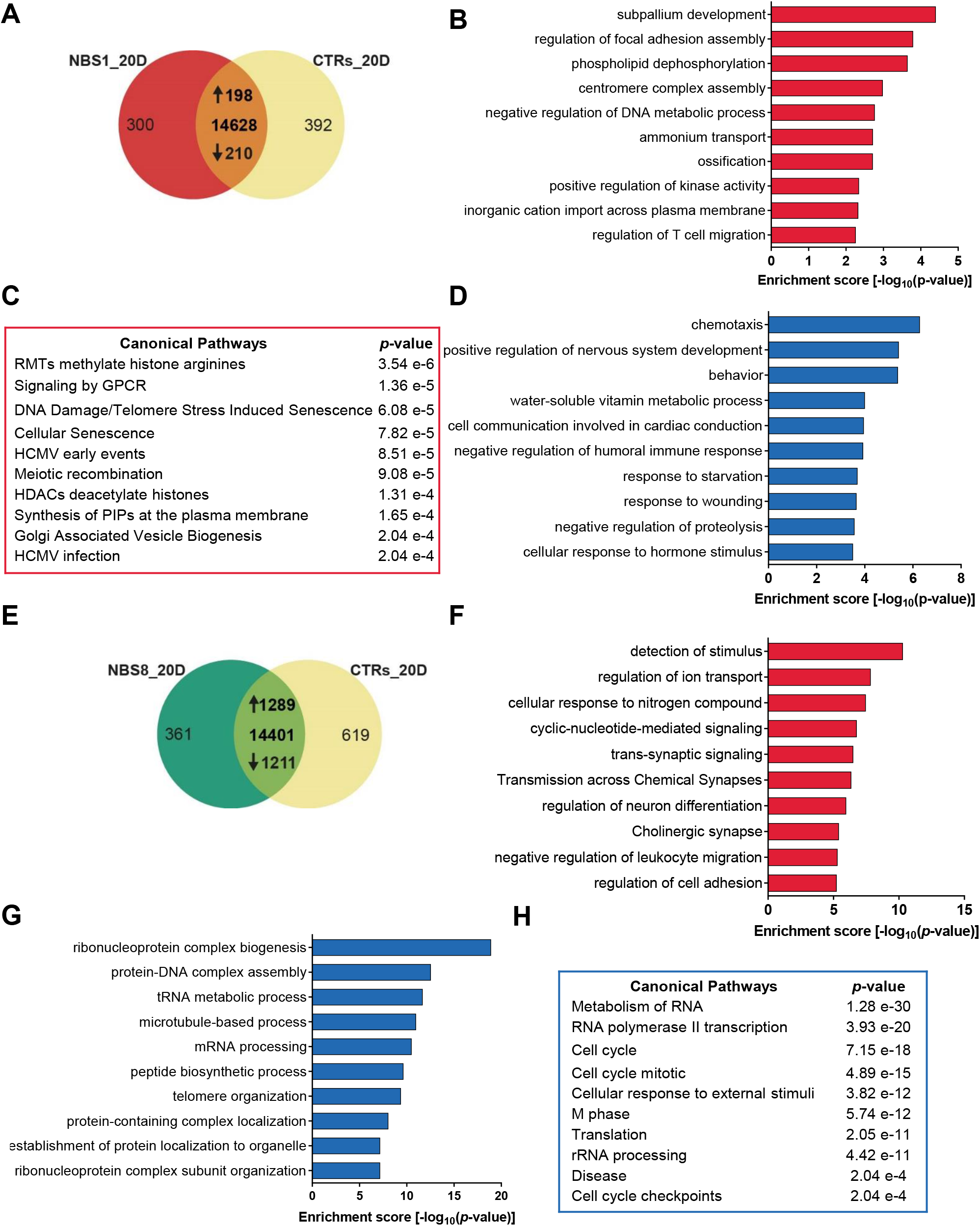
Global transcriptome functional analysis of control and NBS organoids at day 20. **(A)** Venn diagram showing genes expressed only in NBS1 organoids (300), in control organoids (392) and common to both organoids (12828; detection p value < 0.05) **(B)** Bar chart of the enriched clustered GOs (Top 10 ranked) of the significantly up-regulated 198 genes in NBS1-organoids compared to control-organoids. **(C)** Canonical pathways enrichment analysis of the up-regulated 198 genes in NBS1-organoids compared to control-organoids (Top 10 ranked). **(D)** Bar chart of the enriched clustered GOs (Top 10 ranked) of the significantly down-regulated 210 genes in NBS1-organoids compared to control-organoids. **(E)** Venn diagram showing genes expressed only in NBS8-organoids (361), in control-organoids (619) and common to both organoids (14401; detection p value < 0.05) **(F)** Bar chart of the enriched clustered GOs (Top 10 ranked) of the significantly up-regulated 1289 genes in NBS8-organoids compared to control organoids. **(G)** Bar chart of the enriched clustered GOs (Top 10 ranked) of the significantly down-regulated 1211 genes in NBS8-organoids compared to control-organoids. **(H)** Canonical pathways enrichment analysis of the down-regulated 1211 genes in NBS1-organoids compared to control-organoids (Top 10 ranked).

The same analysis was performed using the NBS8-organoids. 1289 genes were identified as up-regulated compared to both control-organoids. **(Figure 3E, Table S2)**. Among the enriched GOs, we could identify clusters connected with synaptic signalling and regulation of neuron differentiation, as well as regulation of ion transport and regulation of cell adhesion **(Figure 3F)**. On the other hand, the cluster analysis of the 1211 down-regulated genes identified *ribonucleoprotein complex biogenesis*, *protein-DNA complex assembly, microtubule-bases process* and *telomere organization* as the most enriched GOs **(Figure 3G)**. The analysis of the canonical pathways identified genes involved in *Metabolism of RNA*, *RNA polymerase II transcription*, *Cell cycle*, *Cell cycle mitotic* and *Cell cycle checkpoints* pathways to be significant reduced **(Figure 3H)**. Inherently, NBS1- and NBS8-organoids show variable modulation of genes associated with regulation of gene expression such *chromatin assembly*, *DNA replication-dependent nucleosome assembly*, *negative regulation of gene expression/epigenetic* and *DNA packing* which were found up-regulated in NBS1 and down-regulated in NBS8 **(Table S2)**. Notwithstanding, both NBS1 and NBS8 organoids downregulated genes associated with behaviour and cognition **(Table S2)**. Our results indicate that although both NBS1- and NBS8-organoids carry the hypomorphic 657del5 mutation, heterozygous and homozygous carriers result in slightly different phenotypes.

### NBS organoids show accumulation of DNA damage

Defective DDR is a well-established cellular feature observed in NBS patients [34]. However, the precise mechanism how NBS cells respond to DSBs and how it affects brain development is poorly understood. In view of this, we took an extensive look into the DNA-damage response at day 20 of our cerebral organoids. At this time-point, we found deregulation in the expression of genes involved in the DDR pathway **(Figure 4A)**. Down-regulation of *ATM* was observed, which was confirmed by qRT-PCR **(Figure 4B)**. Interestingly expression of *TP53* was not significantly regulated **(Figure 4C)**. However, when the protein level was analysed, NBS-organoids clearly showed a reduction in p53 levels compared to control-organoids **(Figure 4D)**.

**Figure 4.**
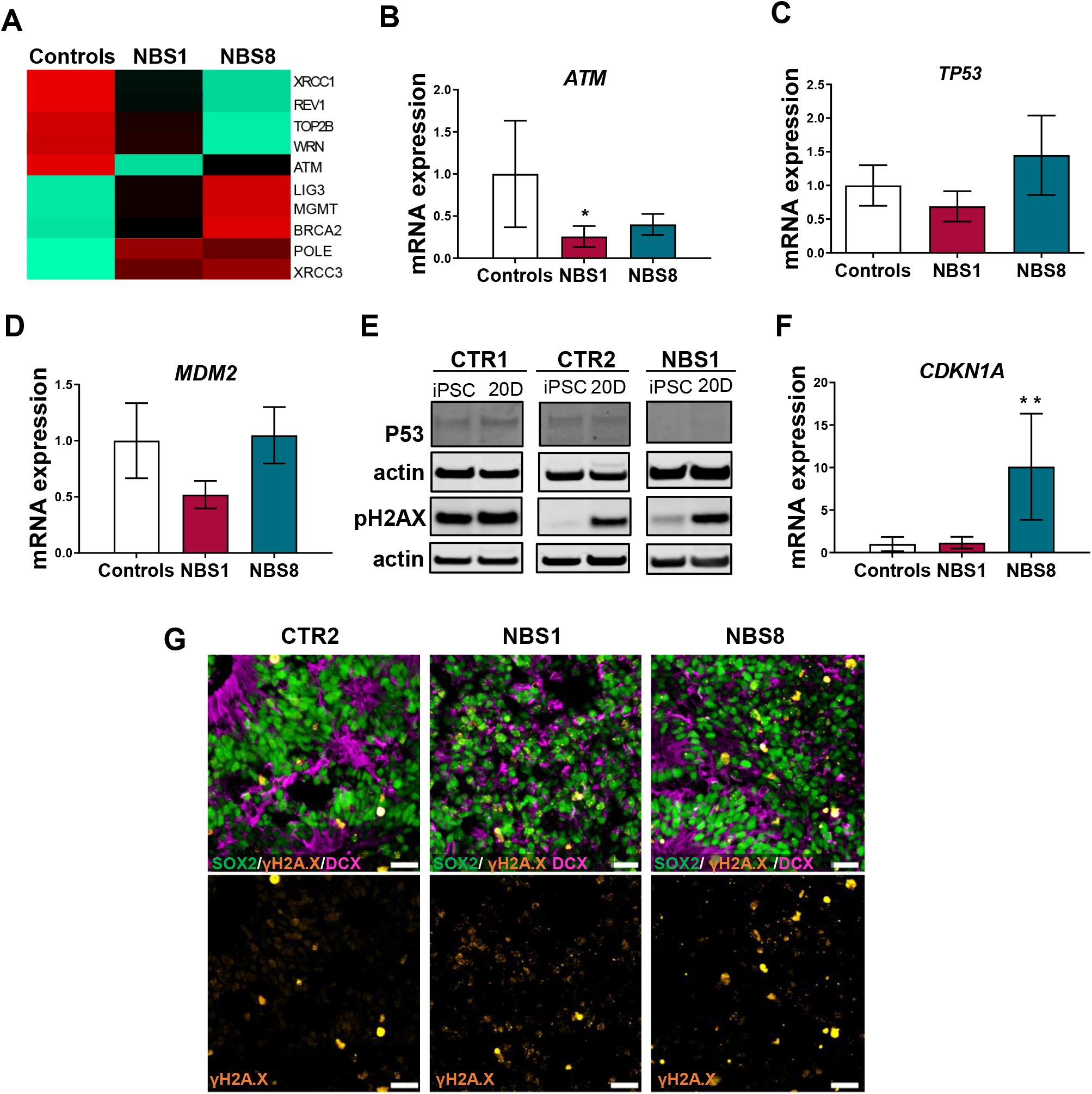
DNA damage response analysis in NBS organoids. **(A)** Heatmap showing differential gene expression analysis of selected DNA damage repair-related genes expressed in control- and NBS-organoids at day 20. **(B-C)** qRT-PCR analysis of *ATM* **(C)** and *TP53* **(D)** mRNA expression in NBS1 and NBS8-organoids relative to control organoids (CTR1 and CTR2). Results are mean ± 95% confidence interval derived from 3 independent differentiations. Significance in comparison to control was calculated with one-way ANOVA followed by Dunnett’s multiple comparison test. *p<0.05 **(D)** Immunoblotting for total p53 and phosphorylated histone H2A.X in CTR1, CTR2 and NBS1 iPSCs and 20 days cerebral organoids. **(E-F)** qRT-PCR analysis of *MDM2* (E) and *CDKN1A* (F) mRNA expression in NBS1 and NBS8-organoids relative to control organoids (CTR1 and CTR2). Results are mean ± 95% confidence interval derived from 3 independent differentiations. Significance in comparison to control was calculated with one-way ANOVA followed by Dunnett’s multiple comparison test. **p<0.01. **(G)** Representative confocal pictures of immunostainings for SOX2, γH2A.X and DCX in CTR2-, NBS1- and NBS8-organoids. NBS-organoids display and increase in γH2A.X nuclear foci formation. Scale bars, 50μm.

To examine if the decrease in p53 expression was due to the presence of mutations, *TP53* target sequencing was carried out in order to identify disease-causing variants. No mutations were found in control- or NBS-organoids (data not shown). As MDM2 is a key regulator of p53, *MDM2* mRNA expression was analysed and no significant d ifferences between control- and NBS-organoids were found **(Figure 4D)**. Surprisingly, NBS8-organoids expressed significantly higher levels of *CDKN1A* **(Figure 4F)**, the major p53 target which controls cell cycle arrest after DNA damage [35]. Outstandingly, high levels of DSBs were observed in the NBS-organoids, judged by the increase of γ-H2AX nuclear foci formation co-localized with the SOX2^+^ NPCs **(Figure 4G).** We next asked whether the observed phenotype could be rescued by stabilization of p53. Thus, cerebral organoids were incubated with Nutlin-3a, an MDM2 inhibitor **(Figure S3A)**. After 72h of incubation (day 25) both CTR1- and NBS8-organoids started to have a dark appearance, indicating cell death **(Figure S3B)**. The inhibition of p53 degradation led to the decrease in the level of *TP53* mRNA in CTR1-organoids, however no differences were observed in NBS8-organoids, were *TP53* levels remained very low **(Figure S3C)**. A similar pattern was observed for *MDM2* expression. Remarkably, *CDKN1A* expression was dramatically up-regulated after Nutlin-3a in both CTR1- and NBS8-organoids **(Figure S3D)**. Overall, Nutlin-3a led to a remarkable decrease in cell proliferation, also confirmed by the down-regulation of *KI67* mRNA expression in both CTR1- and NBS-8 organoids **(Figure S3D)**.

Together, our results indicate that NBS-organoids display an impair DDR pathway, with the accumulation of DNA damage and consequently genomic instability most probably due to the lower levels of p53. Moreover, stabilization of p53-induced by Nutlin-3a led to cell cycle arrest. This approach suggests that p53 plays a central role in orchestrating the fate of NPCs in NBS-organoids.

### NBS organoids exhibit premature differentiation accompanied by NNAT overexpression

To gain deeper insights into the molecular portraits of NBS-regulated gene expression, we analysed the exclusively expressed genes between NBS1- and NBS8-organoids compared to control. We identified 124 genes expressed exclusively in the NBS- and not in the control-organoids **(Figure 5A)**. Functional enrichment analysis revealed that most of these genes are involved in *synaptic signalling*, *presynapse assembly* and *organ induction* **(Figure 5B)**. Besides the genes directly linked with neuronal development, we found genes involved in the *regulation of interleukin-6* production and *matrisome-associated*, including regulators of the ECM and secreted factors. As already described, NBS-organoids showed up-regulation of a number of neuronal differentiation markers **(Figure 1C)**.

**Figure 5.**
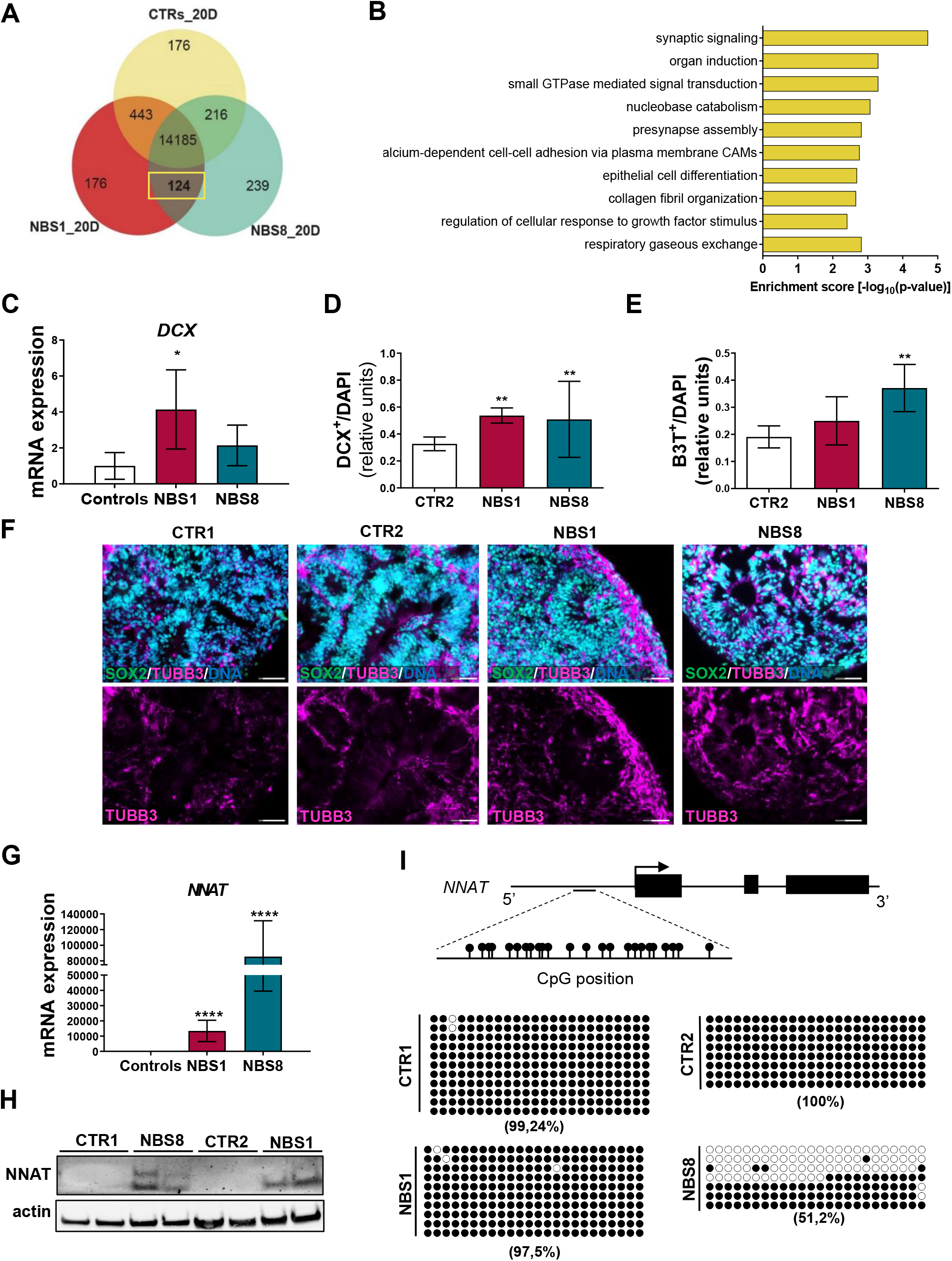
Neurodifferentiation propensity of NBS organoids. **(A)** Venn diagram showing the exclusively expressed genes in NBS1- and NBS8-organoids (124) compared with in control-organoids (detection p value < 0.05). **(B)** Bar chart of the enriched clustered GOs (Top 10 ranked) of the exclusively expressed genes (124) NBS1-and NBS8-organoids compared to control organoids. **(C)** qRT-PCR analysis o*f DCX* mRNA expression in NBS1 and NBS8-organoids relative to control organoids (CTR1 and CTR2). Results are mean ± 95% confidence interval from 3 independent differentiations. **(D-E)** Quantification of the DCX- (D) and βIII-Tubulin (E) positive cells in 20 days CTR2-, NBS1- and NBS8-organoids. Results are mean ± SD from 3 organoids from independent differentiations. **(F)** Representative pictures of immunostainings of SOX2 and βIII-Tubulin in CTR1-CTR2-, NBS1- and NBS8-organoids showing an increase in βIII-Tubulin+ cells. **(G)** qRT-PCR analysis of *NNAT* mRNA expression in NBS1 and NBS8-organoids relative to control organoids (CTR1 and CTR2). Results are mean ± 95% confidence interval from 3 independent differentiations. Significance in comparison to control was calculated with one-way ANOVA followed by Dunnett’s multiple comparison test. ****p<0.0001. **(H)** Immunoblotting for NNAT in CTR1- and NBS8-organoids at day 20 and CTR2- and NBS1-organoids at day 40. **(I)** Bisulfite sequencing of a CpG island within the promotor region of *NNAT*. The % of methylated CpG dinucleotides in CTR1-, CTR2-, NBS1- and NBS8-organoids at day 20 is shown. Filled circles denote methylated CpG dinucleotides. White circles denote unmethylated CpGs.

To evaluate the differentiation propensity of NPCs, we analysed the expression of the neuronal marker DCX. Although mRNA expression was only upregulated in the NBS1-organoids **(Figure 5C)**, both NBS1- and NBS8-organoids showed a significantly higher number of DCX^+^ cells **(Figure 5D)**, thus suggesting premature neuronal differentiation. Likewise, NBS-organoids showed an increase in the number of βIII-Tubulin^+^ cells, but more pronounced in the NBS8-organoids **(Figure 5E,F)**.

Interestingly, we identified *neuronatin* (*NNAT*) as the most up-regulating gene in day 20 NBS-organoids. *NNAT* mRNA expression was barely detectable in the control organoids and highly upregulated in NBS organoids **(Figure 5G)**. Consistent with the mRNA expression, NNAT protein was only present in NBS-organoids **(Figure 5H)**. Regulation of NNAT expression depends on the degree of *NNAT* methylation [36]. Thus, we analysed the methylation status within a CpG island within the promoter region. We found high levels of methylation in this region in the control-organoids (CTR1=99,24% and CTR2=100%) and NBS1-organoids (97,5%). However, a markedly decrease in the methylation at this CpG island was observed in NBS8 organoids (51,2%) **(Figure 5I)**.

Collectively, our results suggest that NBS-organoids undergo premature neurogenesis governed by NNAT and this abnormal over-expression is controlled by the loss of methylation in a regulatory region within the *NNAT* promoter.

### NBS organoids at day 40 acquire an abnormal regulation of cell cycle

To further investigate the consequences of NBS in postmitotic neurons, we differentiated the cerebral organoids for 40 days and performed single organoid transcriptome analysis (SOT-analysis). To ensure the maturity of the organoids, dissociation into single cells for further FACS analysis and re-plating in 2D was performed **(Figure 6A)**. As before, we examined the expression of SOX2, βIII-Tubulin, DCX and KI67. Compared with control-organoids, NBS-organoids were composed of fewer SOX2^+^ cells (NBS=45.97%; control=59.48%). On the other hand, the population of βIII-Tubulin^+^ cells was slightly higher (NBS=35.15%; control=28.6%) **(Figure S3A-C)**, suggesting a more differentiated population in NBS organoids. Around 95% of the NBS-organoids are composed of DCX+ cells, were 20% of these cells are proliferating indicated by the presence of DCX^+^/KI67^+^ cells **(Figure S3A-C)**. These results are in line with previous observations of an increase in DCX expression at day 20.

**Figure 6.**
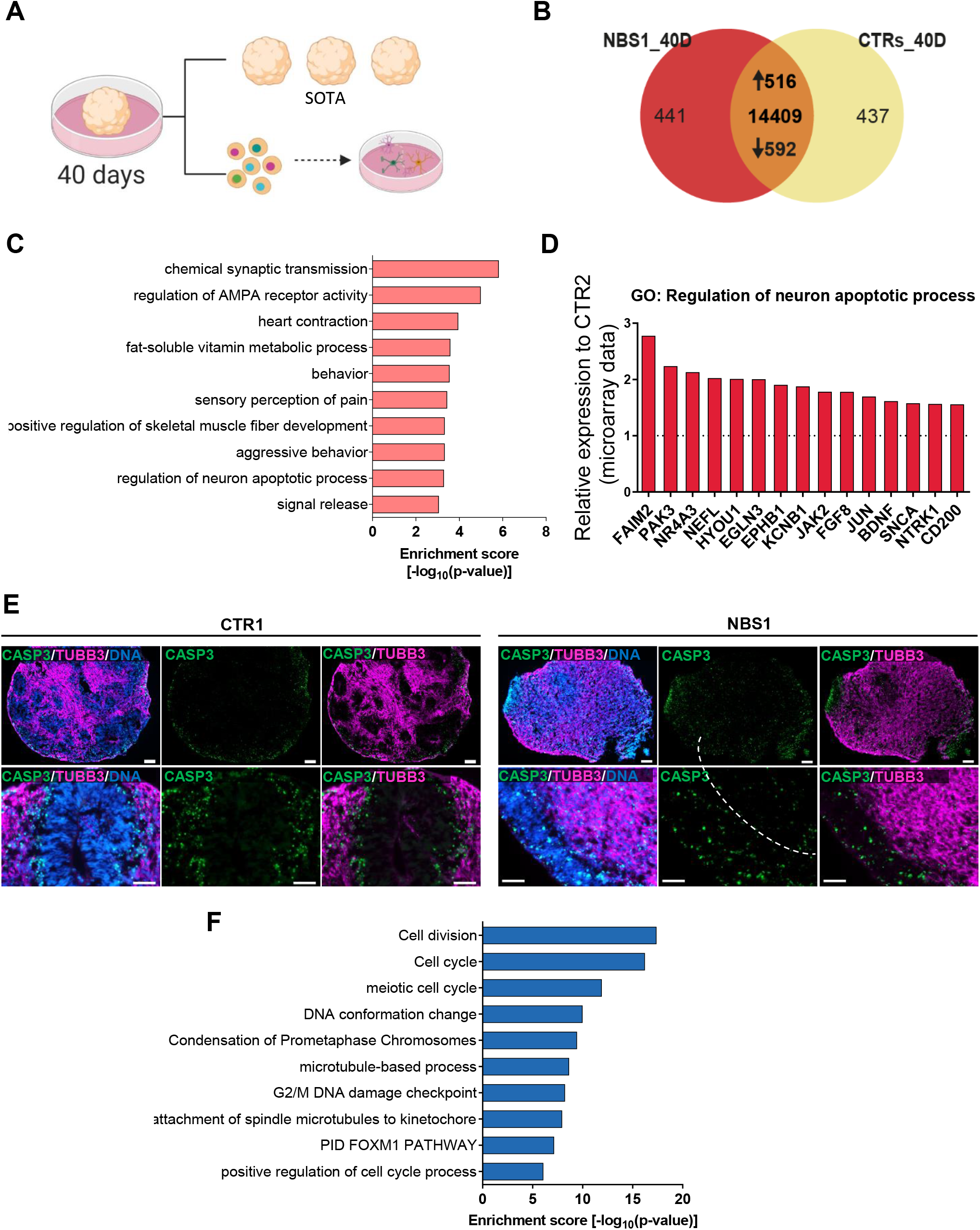
NBS-organoids profile at day 40. **(A)** Schematic depicting the analysis of the NBS and control organoids at day 40. SOT-analysis was performed. Also, cerebral organoids were dissociated into single cells for FACS analysis and re-plating in 2D. SOT: single organoid transcriptome. **(B)** Venn diagram showing genes expressed only in NBS1-organoids (441), in control-organoids (437) and common to both at day 40 (14409; detection p value < 0.05). **(C)** Bar chart of the enriched clustered GOs and Pathways (Top 10 ranked) of the up-regulated genes (1516) in NBS1-organoids compared to control organoids at day 40. ***(D)*** mRNA expression of *FAIM2, PAK3, NR4A3, NEFL, HYOU1, EGLN3, EPHB1, KCNB1, JAK2, FGF8, JUN, BDNF, SNCA, NTRK1* and *CD200* genes part of the GO: Regulation of neuron apoptotic process in NBS1-compared to CTR2-organoids at day 40. Gene expression extracted from the SOT-analysis. **(E)** Representative pictures of immunostainings of cleaved-CASP3 and βIII-Tubulin in CTR1- and NBS1-organoids showing an increase apoptosis in NBS1-organoids. **(F)** Bar chart of the enriched clustered GOs (Top 10 ranked) of the down-regulated genes (592) in NBS1-organoids compared to control-organoids at day 40.

After re-plating in 2D, we could still observe the presence of SOX2+ NPCs and a neuronal network formed by the βIII-Tubulin^+^ neurons **(Figure S3D)**, showing the integrity of our cerebral organoids after 40 days of differentiation.

SOT-analysis identified 516 upregulated and 592 down-regulated genes in NBS1-organoids compared to control-organoids (CTR1 and CTR2) **(Figure 6B, Table S3)**. Analysis of the enriched GOs showed a persistent upregulation of *chemical synaptic transmission* and *regulation of AMPA receptor activity* in the NBS1-organoids **(Figure 6C)**. Interestingly upregulation of *FAIM2*, *PAK3*, *NR4A3*, *NEFL*, *HYOU1*, *EGLN3*, *EPHB1*, *KCNB1*, *JAK2*, *FGF8, JUN*, *BDNF*, *SNCA*, *NTRK1* and *CD200* involved in the regulation of neuron apoptotic process was observed **(Figure 6D).**

We therefore evaluated the presence of apoptosis by immunostaining for cleaved CASP-3 together with the neuronal marker βIII-Tubulin in the cerebral organoids. At day 40, we detected some apoptotic cells in control organoids in the areas where βIII-Tubulin^+^ neurons reside. However, NBS-organoids besides the unorganized cyto-architecture observed with the lack of the VZ, the number of cleaved CASP-3^+^ cells was high. Interestingly, these apoptotic cells were distributed throughout the entire organoid, affecting not only the βIII-Tubulin^+^-neurons but also the NPCs **(Figure 6E)**.

Interestingly, among the downregulated DEGs, transcripts related to *cell division*, *cell cycle*, *DNA G2/M DNA damage checkpoint* were identified **(Figure 6F)**. The transcripts with reduced expression included *CHEK1*, *NBN, MRE11*, *CCNA2*, *CDK1*, *CDC42*, *CCNB1*, *CDKN2C* and *RAD9B.* Furthermore, DNA repair, regulation of TP53 activity were also observed as down-regulated GOs **(Table S3)**.

Taken together, our results imply that NBS-organoids undergo premature neurodifferentiation along with increased apoptosis. Furthermore, SOT-analysis points to a deregulation of cell cycle as the pivotal mechanism underlying perturbed neurodevelopment in the NBS organoids.

### Bleomycin-induced cytotoxicity highlights the aberrant NBS phenotype

To understand the effects of how increased accumulation of DNA damage affects brain development in NBS patients, we subjected the cerebral organoids to bleomycin treatment over a period of 72h and subsequent SOT-analysis. **(Figure 7A)**. Evaluation of DEGs in control organoids after bleomycin treatment revealed an up-regulation of 139 genes **(Figure 7B)**. The subsequent functional enrichment analysis showed that these genes are part of the *P53 downstream pathway* and *TP53 regulates transcription of cell death receptors and ligands* **(Figure 7C)**, thus highlighting bleomycin-induced DNA damage as having a significant effect on the p53 signalling pathway in control organoids.

**Figure 7.**
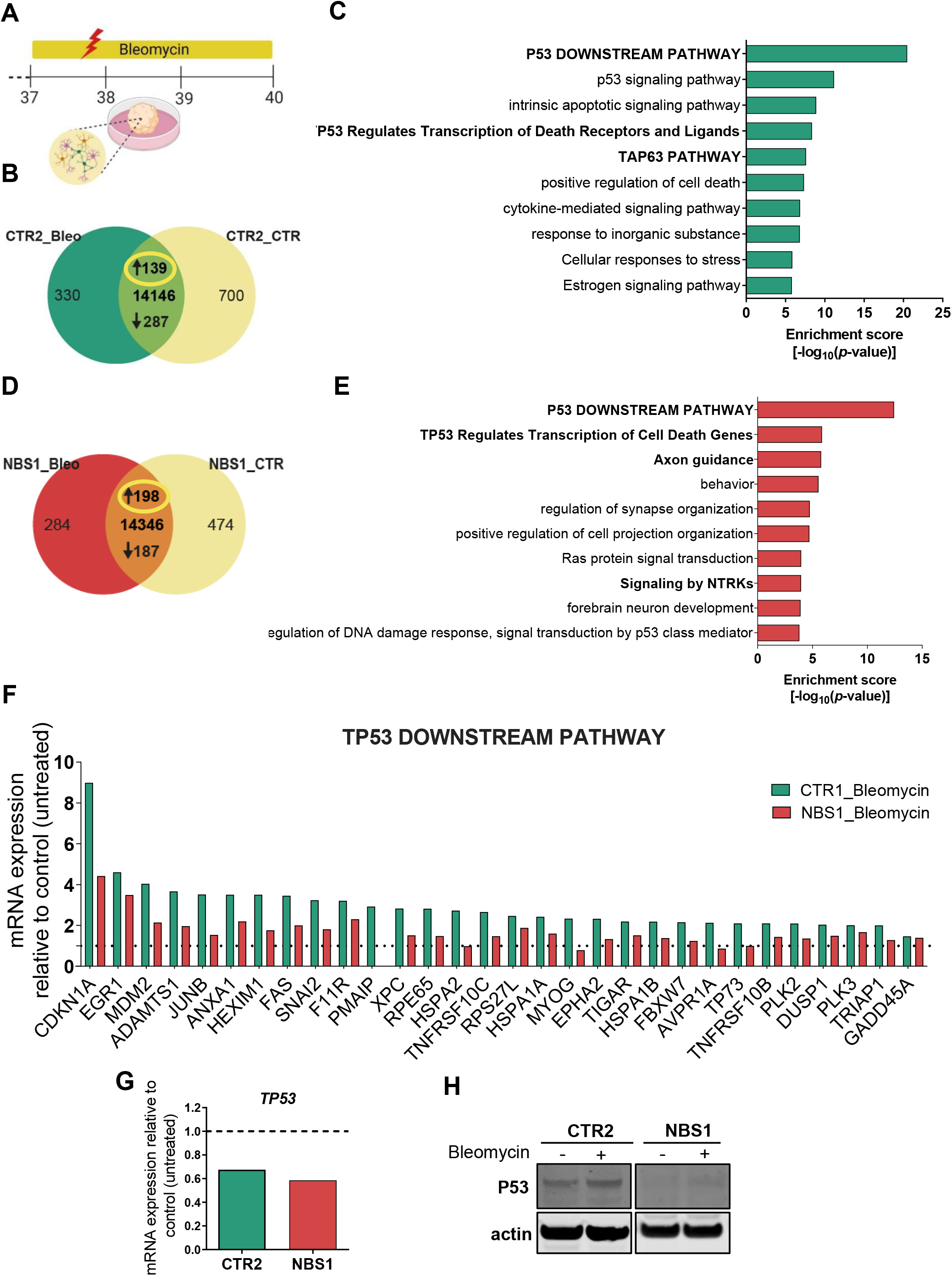
Effects of bleomycin in NBS-organoids at day 40. **(A)** Schematic depicting the strategy to induce DNA damage with Bleomycin treatment during 72h from day 37 to day 40 of CTR2- and NBS1-organoids. **(B)** Venn diagram showing genes expressed only in CTR2_Bleomycin organoids (330), in CTR2_control organoids (700) and common to both (144146; detection p value < 0.05). **(C)** Bar chart of the enriched clustered GOs and Pathways (Top 10 ranked) of the up-regulated genes (139) in CTR2_Bleomycin-organoids compared to CTR2_control-organoids. **(D)** Venn diagram showing genes expressed only in NBS1_Bleomycin-organoids (284), in NBS1_control-organoids (474) and common to both (414346; detection p value < 0.05). **(E)** Bar chart of the enriched clustered GOs and Pathways (Top 10 ranked) of the up-regulated genes (198) in NBS1_Bleomycin-organoids compared to NBS1_control-organoids. **(F)** mRNA expression of the genes part of the up-regulated *TP53 downstream pathway* in CTR2_Bleomycin-organoids and NBS1_Bleomycin-organoids compared to CTR2_control-organoids and NBS1_control-organoids, respectively. All genes were significantly up-regulated in CTR2 after bleomycin treatment. **(G)** mRNA expression of *TP53* in CTR2_Bleomycin-organoids and NBS1_Bleomycin-organoids compared to CTR2_control-organoids and NBS1_control-organoids. **(H)** Western blot analyses of total p53 in CTR2- and NBS1-organoids after bleomycin treatment.

Focusing on NBS organoids, 198 up-regulated genes were identified when comparing bleomycin treatment to control conditions **(Figure 7D)**. Interestingly, these genes were also enriched for *P53 downstream pathway-TP53 regulates transcription of cell death genes*. Furthermore, bleomycin treatment re-inforces axon guidance, regulation of synaptic organization and forebrain development, already exacerbated in basal conditions in the NBS-organoids **(Figure 7E).**

We next evaluated the levels of mRNA expression of the genes up-regulated within the *TP53 downstream pathway* between control and NBS organoids after bleomycin treatment. While both CTR2- and NBS1-organoids activate the p53 signalling pathway to mediate DDR, NBS organoids showed an impairment in the activation of these genes compared to CTR2-organoids, as observed by the significantly lower expression of these genes which include *CDKN1A, FAS* and *GADD45A* **(Figure 7F)**.

Additionally, the analysis of the DEGs upon bleomycin treatment showed 287 down-regulated genes in control organoids **(Figure S5A)**. These genes were highly significantly enriched for cell cycle and cell cycle checkpoints (Figure S5B). NBS1-organoids followed the same pattern, with 187 down-regulated genes **(Figure S5C)** being enriched for *cell cycle*, *cell division*, *Mitotic G1 phase and G1/M transition* and *DNA Double-strand break repair* **(Figure S5D).** Next, we evaluated the expression of the common set of genes between CTR2 and NBS1-organoids which belong to the *cell cycle pathway*. While CTR2-organoids significantly down-regulated cell cycle-related genes, NBS1 organoids were not able to induce such dramatic changes in gene expression after bleomycin treatment **(Figure S5E)**.

The consequences of these impairments are correlated with the expression of *MKI67,* where CTR2-organoids drastically reduced *MKI67* levels in contrast with a slight reduction observed in NBS1-organoids **(Figure S5F)**. To have a better understanding of the specific effects of bleomycin treatment in NBS organoids, we performed enrichment analysis of the 94 exclusively expressed genes **(Figure S5G).** Bleomycin treatment specifically induced the expression of genes associated with *trans-synaptic signalling*, *glycosphingolipid biosynthesis*, *leukocyte apoptotic process* and *neurotransmitter transport* in the NBS-organoids **(Figure S5H)**.

Taken together, our results have shown that after exposure to a genotoxic agent, NBS organoids undergo an impairment in P53-regulated pathways and consequently cell cycle regulation due to the low levels of p53.

## Discussion

While microcephaly is the hallmark of NBS, the mechanisms that lead to reduced brain size in these patients are largely unknown mostly due to the hurdles and limitations associated with studying NBS. By generating for the first-time patient iPSC-derived cerebral organoids we were able to investigate cellular and molecular effects of the *NBN* mutation during early neurogenesis.

In this work, two distinct NBS patient-derived iPSC lines carrying the hypomorphic *NBN* 657del5 mutation were analysed-NBS1 and NBS8. NBS is an autosomal recessive disorder however NBS8-iPSCs are heterozygous for the *NBN* 657del5 mutation, indicating that other missense mutations within *NBN* could be present [37]. NBS8-iPSCs present a much higher chromosomal instability, with aberrations acquired during the reprogramming process, as such duplications of chromosome 5q, which can confer growth advantages during the reprogramming process [38].These differences in the genotype of both iPSC lines can result in distinct phenotypes. Indeed, a variable phenotype as a result of neurodevelopmental abnormalities has been observed in a number of NBS patients [7].

NBS-organoids has allowed not only to identify distinct phenotypes but also to identify common mechanisms underlying the etiology of brain abnormalities in NBS patients. As NPCs and post-mitotic neurons respond differently to endogenous DNA damage [1], we analysed the cerebral organoids at two time-points: 20 days and 40 days of differentiation. Day 20 cerebral organoids correspond to 8-9 pcw in human brain development and are composed mainly by cells with a forebrain identity. However, a sub-population of cells with midbrain and hindbrain identity were also present as observed by others [24]. Microcephaly-associated abnormalities are a hallmark of NBS which frequently occur during the early neurodevelopmental stages, although in some patients only develop postnatally [7,39]. While both NBS1- and NBS8-organoids showed a similar expression profile for the analysed differentiation markers, morphology-based analysis revealed distinct phenotypes. Although an increase in size at day 20 was observed, NBS8-organoids (heterozygous) presented NPCs aligned in the apical membrane of the VZs and neurons in the cortical region similar to control-organoids. On the other hand, NBS1-organoids (homozygous) were significantly smaller in size and presented a disrupted architecture with a disorganized distribution of cells, resulting in less and small VZs, thus recapitulating the microcephaly phenotype. As already mentioned, the different morphologies between the NBS-organoids can be attributed to the distinct genetic composition and highlights the potential of using cerebral organoids to model NBS.

So far, the use of brain organoids to model microcephaly has only been performed with iPSC-derived from patients carrying centrosome-related mutations. Apart from the smaller size of those organoids, a depletion of NPCs was observed [24,40–42]. Interestingly, neither the number nor the proliferation of NPCs was affected in NBS-organoids in comparison to the controls. These results hint at a different mechanism underlying microcephaly in NBS patients.

Transcriptome-based analysis after 20 days of differentiation reinforced that NBS1- and NBS8-organoids respond in distinct manner to the endogenous levels of oxidative stress due to the proliferation of the NPCs. NBS1-oganoids showed up-regulation of genes involved in epigenetic regulation of gene expression, centromere complex assembly and senescence, implying these cells attempt to maintain genomic and epigenomic integrity, since DNA methylation and chromatin remodelling are linked to DNA damage and repair [43]. However, a number of genes that regulate gene expression were found down-regulated in the NBS8-organoids, together with genes associated with cell cycle and cell cycle check-points.

Following DSBs, ATM is activated by auto-phosphorylation and undergoes spatial relocation to the DSB site, followed by the phosphorylation of H2AX. In turn, H2AX recruits the MNR complex to the DSB, a process which requires NBN. The DNA damage response is then reinforced by further deposition of ATM at the DSB site, promoted by the MNR complex. Besides sensing DSBs, NBN phosphorylates ATM to control cell cycle checkpoints [44,45]. Among the several genes within the DDR pathway de-regulated in the NBS-organoids, *ATM* was found significantly down-regulated, in accord with previous studies demonstrating that NBS cells dramatically reduce ATM activation [46]. The ATM-dependent signaling events lead to rapid p53 stabilization and transcriptional induction of *CDKN1A*, which encodes p21- a cyclin-dependent kinase inhibitor which triggers G1 cell cycle arrest or a permanent state of senescence or apoptosis [44,45]. NBS-organoids showed a dramatic reduction in p53 protein levels, with no differences in *TP53* mRNA expression. Our results are in line with our previous study where we showed that NPCs derived from NBS-iPSCs express low levels of p53 [23]. In an early neurodifferentiation phase, p53 is still able to activate the transcription of *CDKN1A* (p21) in the NBS-organoids, thus demonstrating that p53 function is not completely abrogated. However, the DDR pathway is compromised, as shown by an increase in γ-H2AX nuclear foci formation in the NPCs of the NBS-organoids, thus higher levels of DNA damage.

As an attempt to increase the stabilization of p53, we treated the cerebral organoids with nutlin-3a, which functions as to inhibit the interaction between MDM2 and p53 [47]. NBS-organoids underwent significant up-regulation of *CDKN1A*, leading to a down-regulation of *MKI67* and subsequent cell death. Our results indicate that inhibition of MDM2-p53 interaction by nutlin-3a allows p53 stabilization and the enhanced response to nultin-3a further demonstrates that the levels of DNA damage are higher in NBS-organoids. Our data supports previous findings showing that the presence of the truncated p70-nibrin, as a result of the 647del5 *NBN* mutation, is able to retain residual activity to ensure survival [10]. However, an impaired DDR pathway was observed in the NBS-organoids, probably due to the attenuation of the ATM-p53 pathway activation after endogenous DNA damage, in line with previous studies [48,49]. Our data suggests this impairment has an impact on the fate of the NPCs, leading to premature differentiation.

NBS-organoids exclusively expressed genes associated with synaptic signalling and neuronal differentiation as indicated by the increased number of DCX and βIII-tubulin^+^ cells. Our results are in contrast with the previous findings in NBS using NPCs derived from NBS-iPSCs showing delayed neurogenesis [23]. This discrepancy could be due to the different approaches used, such as different differentiation protocols and the iPSC were differentiated only to the NPC stage in Halevy *et al.* (2016).

Premature differentiation of the NPCs leading to its depletion is the common mechanism underlying microcephaly described in patient-derived organoids [24,40–42]. The accumulation of DNA-damage can promote accelerated neurogenesis by increasing the expression of genes that regulate cellular, circuity and cognitive functions which includes neurite outgrowth and synapsis development, as our data shows [50]. During neurodevelopment, DNA methylation is a key process that regulates the expression of these genes and therefore the maintenance and fate specifications of NPCs [51]. Our results suggest that the premature differentiation observed could be explained by the over-expression of NNAT as a result of the loss of methylation within a CpG island in the *NNAT* promoter observed in the NBS8-organoids. Despite the methylation profile of NBS1-organoids was similar to both control organoids, we cannot entirely exclude the presence of hypomethylation in another regulatory region of the *NNAT* promoter. Hence, further detailed methylation analysis in the NBS-organoids should be performed. NNAT is a small transmembrane proteolipid that has been reported to promote neuronal differentiation and a key molecule that maintains cellular homeostasis [52,53]. Interestingly, NNAT expression has been associated with the regulation of stress levels in cells. However, in addition to its protective role, over-expression of NNAT is frequently found in patients with glioblastoma and contributes to the neuronal pathogenesis of in Lafora disease due its strong propensity to misfold and aggregate leading to subsequent apoptosis and neuronal loss [53,54]. Day 40 transcriptomic data revealed an increase in neuron apoptosis, in contrast no differences in apoptosis was observed in day 20 NPCs. Along with increased apoptosis, NBS-organoids down-regulated genes involved in cell division, cell cycle, chromosome segregation and the G2/M damage checkpoint. The defects in this core pathways observed at day 40 led to the increase in the numerical and structural alterations, both common features of NBS, hence suggestive of a progressive phenotype. With the increase of DNA-damage, mimicked here by bleomycin, NBS-organoids showed delayed activation of p53 downstream targets and as a result not being able to activate DNA damage checkpoints and stop cell division at the same magnitude seen in control organoids.

Together, our data suggests the *NBN* 657del5 mutation induces a progressive phenotype. In an early developmental stage, cells try to maintain homeostasis and genomic integrity. However, due to the inability to efficiently repair damaged DNA, NPCs undergo premature differentiation with disrupted structural organization. We hypothesize that this process is in part triggered by the over-expression of NNAT. The observed increase in genomic instability is a consequence of delayed p53-mediated DNA damage response. Ultimately, the increase in DNA damage together with an aberrant trans-synaptic signalling leads to neuronal apoptosis. We suggest that the premature differentiation of the NPCs and an increase in apoptosis in the NPCs and post-mitotic neurons as the mechanism underlying progressive microcephaly-the hallmark of NBS.

## Materials and Methods

### iPSC derivation and culture methods

The iPSC lines derived from NBS patients (NBS1 and NBS8) as well as the control individuals CTR1 and CTR2 used in this study have been described [21,28,29,55]. iPSC lines were cultured in mTESR medium (Stem Cell Technologies) supplemented with Penicillin/ Streptomycin (P/S) on Matrigel-coated plates (Corning). The medium was changed every day and cells were passaged every 5-6 days using PBS without Calcium and Magnesium (Life Technologies).

### Generation of cerebral organoids

For the generation of cerebral organoids, iPSCs were differentiated as described [56], but with further optimization. On day 0, iPSCs were dissociated using TrypLE Express (Gibco) and plated into a 96-well ultra-low-attachment (10000 cells/well, NucleonTM SpheraTM, Thermo Scientific) in mTEST supplemented with 10 μM ROCK inhibitor Y-27632 (Tocris Bioscience). Neural induction was initiated on day 1 by adding Neural induction medium (NiM) (DMEM/F12, 20% Knock-out serum replacement, 1%NEAA, 0.5% GlutaMAX and 0.1mM 2-Mercaptoethanol (all from Gibco) with 10 μM SB-431542 (Tocris Bioscience), 5 μM Dorsomorphine (Tocris Bioscience) and 10 μM ROCK inhibitor Y-27632. Medium was changed daily. After 5 days the spheroids were transferred to Neural differentiation medium (NdM) (Neurobasal, 2% B27, 1% GlutaMAX, 1% P/S (all from Gibco) supplemented with 20 ng/mL of EGF and FGF2 (both PrepoTech) in non-adhesive 100 mm dishes. Organoids were further cultured under continuous agitation (60 rpm) in a shaker incubator(New Brunswick S41i, Eppendorf) with daily medium change until day 16, and from day 16 onwards medium was changed every other day. At day 25 medium was replaced to NdM supplemented with 20 ng/mL of BDNF with medium change every 2-3 days. From day 40 onwards, organoids were kept in NdM with medium change every 4 days. To induce DNA damage, cerebral organoids at day 37 were incubated with 30 μg/mL of Bleomycin (Millipore) for 72h before harvesting for RNA extraction and for protein lysate preparation.

### Single-cell dissociation and 2D neuronal culture

40-day CTR1-, NBS1- and NBS8-organoids were dissociated to a single cell suspension for 30 minutes using the Papain dissociation kit (Worthington) according to the manufactures’ instructions. 300000 cells were replated into poly-L-ornithine and laminin (Sigma) coated coverslips in neural differentiation medium [(Neurobasal, 1% NEAA, 1% N2, 1% P/S (all Thermo Fisher Scientific) supplemented with 1μM of cAMP (Thermo Fisher Scientific, Rockford, IL, USA) and 10 ng/mL of BDNF, GDNF and IGF-1 (all Immuno Tools)].

### Organoid sectioning and immunostainings

Cerebral organoids were fixed in 4 % paraformaldehyde (PFA) for 30 min at room temperature, washed with PBS and dehydrated in 30% sucrose in PBS overnight at 4°C. Subsequently organoids were transferred into embedding medium (Tissue-Tek OCT Compound 4583, Sakura Finetek), snap-frozen on dry ice and stored at −80°C. Organoids were cut into 10 μm sections and captured in Superfrost Plus slides (Thermo Scientific) using a Cryostat Leica CM3050 S. Cryosections were permeabilized with 0.1% Triton X-100 for 10 min and blocked with 3% BSA in PBS for 1h. Samples were then incubated with the following primary antibodies overnight at 4°C: mouse anti-βIII-tubulin (1:200, CST TU-20), mouse anti-SOX2 (1:100, Invitrogen, Thermo Fisher Scientific 20G5), rabbit anti-SOX2 (1:200, C ST 3579S), guinea pig anti-DCX (1:200, SySy 326004), mouse anti-KI67 (1:200, CST 9449), rabbit anti-cleaved CASP3 (1:200, CST 9664). After washing with PBS, cells were then incubated with the appropriate secondary antibody conjugated with Alexa-488, Alexa-555 or Alexa-647 (1:500, Invitrogene, Thermo Fisher Scientific) for 1h at RT. The nuclear stain Hoechst 33258 (2ug/mL, Sigma) was added at the time of the secondary antibody incubation. Slices were mounted with Fluoromount-G (SouthernBiotech) and fluorescent and confocal images were obtained using a LSM 700 microscope (Carl Zeiss), and processed using ZEN software (Carl Zeiss).

### Quantitative assessment of cerebral organoids and image analysis of histological sections

The size of the organoids was measured employing a Digital Microscope Leica DMS100. Two perpendicular measurements of the organoids diameter (μM) was determined using ImageJ and the mean was calculated. For quantification of the number of ventricular zones-like structures per organoid, organoid slides were stained with SOX2, DCX and nuclei stained with Hoeschst. For quantification of the loop diameter (μM) for each ventricular zones-like structure, two measurements (0 and 45 degrees) were performed, from the apical membrane to the basal membrane, and the mean calculated. To determine the number was SOX2+, KI67+, DCX+, and βIII-Tubulin+ cells relative to the total number of cells, the total integrated intensity for each antibody was calculated and divided by the integrated intensity of the nuclei staining. At least 4 sections per organoid were analysed. Loop diameter, apical membrane length and integrity intensity were measured using ImageJ software.

### Reverse transcriptase PCR (RT-PCR)

Total RNA was extracted from day 20 (3 pooled-organoids) or day 40 (single organoids) using TRIzol (Life technologies) and Direct-zol RNA Mini Prep (Zymo Research) according to the manufacturer’s protocol. 500 ng of purified RNA was used for cDNA synthesis using TaqMan reverse transcription reagent (Applied Biosystems). cDNA was used for subsequent PCR analysis. To detect the presence of transcripts, RT-PCR reactions were performed using 20 ng of cDNA of WT2 and NBS8 organoids using GoTaq® DNA Polymerase (Promega). Total RNA from commercially human fetal and adult brain (BioChain®) was used as positive control. Transcripts abundance was determined by reverse transcription quantitative PCR (qRT-PCR) using the SYBR® Green RT-PCR assay (Applied Biosystems). Amplification, detection of specific gene products and quantitative analysis were performed using a ‘ViiA7’ sequence detection system (Applied Biosystem). The expression levels were normalized relative to the expression of the housekeeping gene RPS16 using the comparative Ct-method 2-Ct. Experiments were carried out in biological triplicates except for 40d organoids (technical triplicates). RT-qPCR data are depicted as mean values with 95% confidence interval. Primers are listed in Table S5.

### Microarray analysis

100 ng of total RNA was subjected to hybridization on the Human Clarion S Array (Affymetrix, Thermo Fisher Scientific) at the BMFZ (Biomedizinisches Forschungszentrum) core facility of the Heinrich-Heine Universität, Düsseldorf. RNA integrity was evaluated using a Fragment Analyzer (Advanced Analytical Technologies, AATI). Data analysis of the Affymetrix raw data was performed in the R/Bioconductor [57] environment using the package affy [58]. The obtained data were background-corrected and normalized by employing the Robust Multi-array Average (RMA) method from the package affy. Hierarchical clustering dendrograms and heatmaps were generated using the heatmap.2 function from the gplots package with Pearson correlation as similarity measure and colour scaling per genes [59]. Expressed genes were compared in venn diagrams employing package VennDiagram [60]. Gene expression was assessed with a threshold of 0.05 for the detection-p-value which was calculated as described in Graffmann et al. [61]. Comprehensive functional analysis of the clustered GO biological processes and pathways (KEGG pathways, Reactome Gene Sets, Canonical pathways and CO-RUM) of the candidate genes was performed using Metascape tool (http://metascape.org) [62]. The default parameters were used: terms with p<0.01, minimum overlap 3 and enrichment factor >1.5.

### Western Blot

Cells of 5 pooled organoids were washed with PBS and then lysed in RIPA buffer containing complete protease and phosphatase inhibitor cocktail (Roche). Lysates were cleared by centrifugation at 20.000g for 10min and quantified with the Pierce™ BCA Protein Assay kit (Thermo Scientific). 25 μg of the lysates were then separated on NuPAGE 4-12% Bis-Tris gels (Invitrogen) and blotted to a 0.45 μm nitrocellulose membrane for 3h at 300mA. The blots were blocked in PBS/ 0,05%Tween20 containing 5% skim milk and then probed with the following primary antibodies over night at 4°C: mouse anti-P53 (1:100, Merck), rabbit anti-H2AX (1:1000, Cell Signaling), mouse anti β-actin (1:4000, Cell Signaling). After washing the blots three times with PBS/ 0,05%Tween20 they were incubated with the appropriate secondary antibody: goat anti-mouse IRDye 680RD and 800CW as well as goat anti-rabbit IRDye 680RD and 800CW (all from LI-COR Biosciences). Following three times washing with TBS/0.05% Tween20 the fluorescent signals were quantified by applying the Odyssey infrared imaging system (LI-COR Biosciences).

### Flow cytometry

The cell marker expression of the 40-day organoids was analyzed by Flow cytometry analysis. To conduct this analysis, 100000 single-cells derived from CTR1-, NBS1- and NBS8-organoids were used. Cell pellet was fixed for 20 minutes in the dark in fixation buffer (Biolegend) followed by permeabilization with permeabilization buffer (Biolegend). Pelleted cells by centrifugation for 5 min ad 300g were stained for 20 min in the dark with the fellowing antibodies: mouse anti-SOX2 (1:200, CST 3579S) mouse anti-βIII-tubulin (1:200, CST TU-20), mouse anti-KI67 (1:200, CST 9449) and guinea pig anti-DCX (1:200, SySy 326004). The cell pellet was washed with permeabilization buffer and incubated with the appropriate secondary antibody Alexa488 and Alexa647 (1:500, Thermo Fisher Scientific). Alexa488 and Alexa647-coupled IgG were used as negative control. Cell pellet was resuspended in PBS/2mM EDTA/0.5%BSA and cell fluorescence was measured using CytoFLEX S (Beckman Coulter). Flow cytometry data were analyzed using FlowJo X v10.6.1 software (FlowJo LLC, Ashland, OR).

### Karyotyping and array comparative genomic hybridization (array-CGH)

The karyotype analysis of NBS-iPSCs was carried out by the Institute of Human Genetics and Antropology, Heinrich-Heine-University, Düsseldorf, Germany. Genomic DNA from NBS- and NBS8-iPSC lines was extracted using the DNeasy Blood Tissue Kit (Qiagen) and array-CGH was performed using the Illumina HumanOmni2.5Exome-8 Bead-Chip v1.3chip at LifeBrain GmbH, Bonn. Genotype and copy number variation (CNV) analysis was performed using Illumina GenomeStudio V2.0.2 (Illumina).

### Analysis of mutational status of *TP53*: Library preparation and massive parallel sequencing

DNA was quantified by a custom-made qPCR assay (Primer for: 5‘ AAACGCCAATCCTGAGTGTC-3‘; Primer rev: 5‘ CATAGCTCCTCCGATTCCAT-3‘). Library preparation was carried out using Ion AmpliSeq™ Library Kit 2.0 and Ion AmpliSeq™ Colon and Lung Cancer ResearchPanel v2 with 10 ng of amplifiable DNA following manufacturer’s recommendations. Ion Xpress™ Barcode Adapters Kits were utilized for barcoding the libraries. Afterwards, libraries were quantified by qPCR using Ion Library TaqMan™ Quantitation Kit on a StepOnePlus™ Real-Time PCR System and were compiled equimolarly for subsequent sequencing reaction. Massive parallel sequencing was conducted on an Ion S5 System using the Ion 520™ Ion 530™ Kit-OT2 with an Ion 530™ Chip. Primary data analyses were performed by Ion Torrent Suite Software. For variant annotation generated Binary Alignment Map (BAM), files were uploaded to and analyzed by Ion Reporter™ Software using recommended analysis parameter for the Ion AmpliSeq™ Colon and Lung Cancer ResearchPanel v2. Detected variants were examined using the Integrative Genomics Viewer (IGV) [63,64]. All reagents and software were from Thermofisher (Darmstadt, Germany). Selected parts of Exon 2, Exon 4 - 8 and Exon 10 from the TP53 gene (NM_000546.5) are covered by the Ion AmpliSeq™ Colon and Lung Cancer ResearchPanel v2 including following amino acids: Ex 2: Met1 - Ser20; Ex 4: Glu68 - Gly112; Ex 5: Tyr126 - Ala138 Ser149 - Gly187; Ex 6: Gly187 - Pro223; Ex 7: Val225 - Glu258; Ex 8: Asn263 - Ala 307; Ex 10: Ile332-Ser367.

### Bisulfite genomic sequencing

Bisulfite conversion of 500ng of DNA of CTR1, CTR2, NBS1 and NBS8-organoids at day 20 was conducted using the EpiTec Kit (Qiagen, Hilden, Germany) as described [65]. PCR primers for specific amplification of NNAT (NG_009263.1) promotor are listed in Table S5. The amplification conditions were denaturation at 95°C for 13min, followed by 35 cycles of 95°C for 60s, 51°C for 50s, and 72°C for 25s. PCR reactions were performed using 25ng bisulfite converted DNA using GoTaq® DNA Polymerase (Promega). The TA Cloning Kit (Invitrogen) was used for cloning of the amplification product (281bp) according to the manufacturer’s instructions. Sanger sequencing was performed at the BMFZ (Biomedizinisches Forschungszentrum) core facility of the Heinrich-Heine Universität, Düsseldorf. 12 clones were sequenced to obtain the methylation profile per sample. Analysis of methylated CpGs and methylation graphs were obtained by using QUMA(http://quma.cdb.riken.jp/) [66] software.

### Statistical analysis

Statistical analysis was performed with GraphPad Prism Software version 8.02 (GraphPad software, San Diego, CA, USA). Ordinary one-way ANOVA was used for statistical significance analysis for comparisons of the mean among groups, followed by a post hoc test with the use of Dunnett‘s multiple comparison test. Statistical significance was assumed at p< 0.05, **p<0.01 and ***p<0.001 and **** p<0.0001. All data are expressed as mean ± 95% confidence interval (qRT-PCR data) or mean ± standard deviation (SD). N and p-values are reported in each figure legend.

## AUTHOR CONTRIBUTIONS

Conceptualization: S.M. and J.A.; Methodology: S.M., L.E., W.G., A.D.; Formal analysis: S.M., W.W., J.A.; Data curation: W.W.; Investigation: S.M.; Resources: K.C.; Writing—original draft preparation: S.M.; Writing—review and editing: J.A.; Supervision: J.A.

## ACKNOWLEDGEMENTS

The authors thank Prof. Gerhard Fritz and Dr. Nina Graffmann for inspiring discussions and comments on the work and the manuscript presented here. Thanks to Martina Bohndorf for technical support.JA acknowledges support from the Medical Faculty, Heinrich-Heine-University, Düsseldorf. This preprint is formatted using a LATEX class by Ricardo Henriques.

## Supplementary material

### Supplementary Tables

**Table S1.**
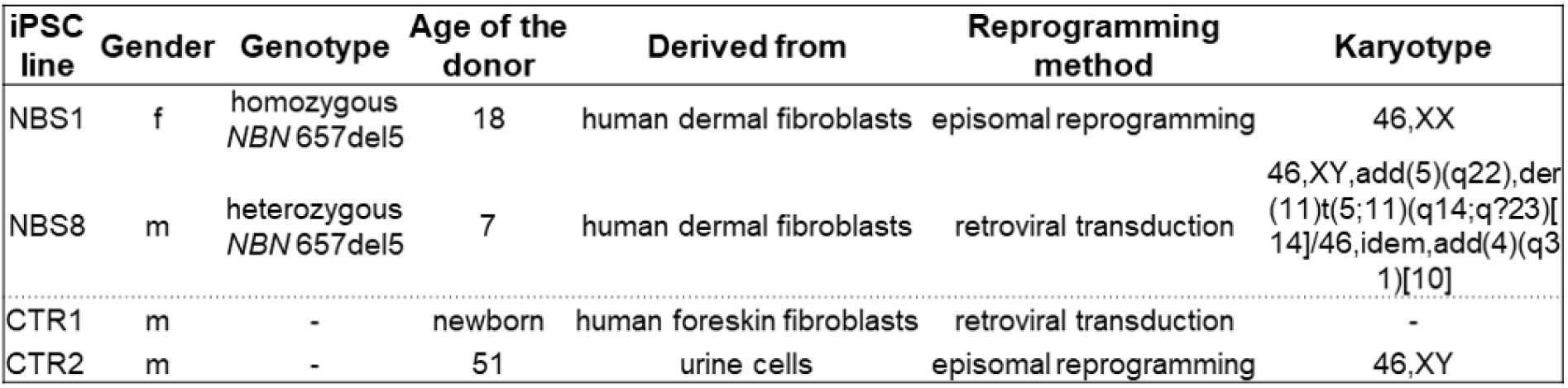
Overview of the iPSC lines used in this study.

**Table S2.** Complete GOs and canonical pathways enrichment analysis of NBS-organoids at day 20 as shown in Figure 3 and 5.

**Table S3.** Complete GOs and canonical pathways enrichment analysis of CTR2- and NBS1-organoids at day 40 as shown in Figure 6.

**Table S4.** Complete GOs and canonical pathways enrichment analysis of the bleomycin treatment in CTR- and NBS-organoids at day 40 as shown in Figure 6.

**Table S5.**
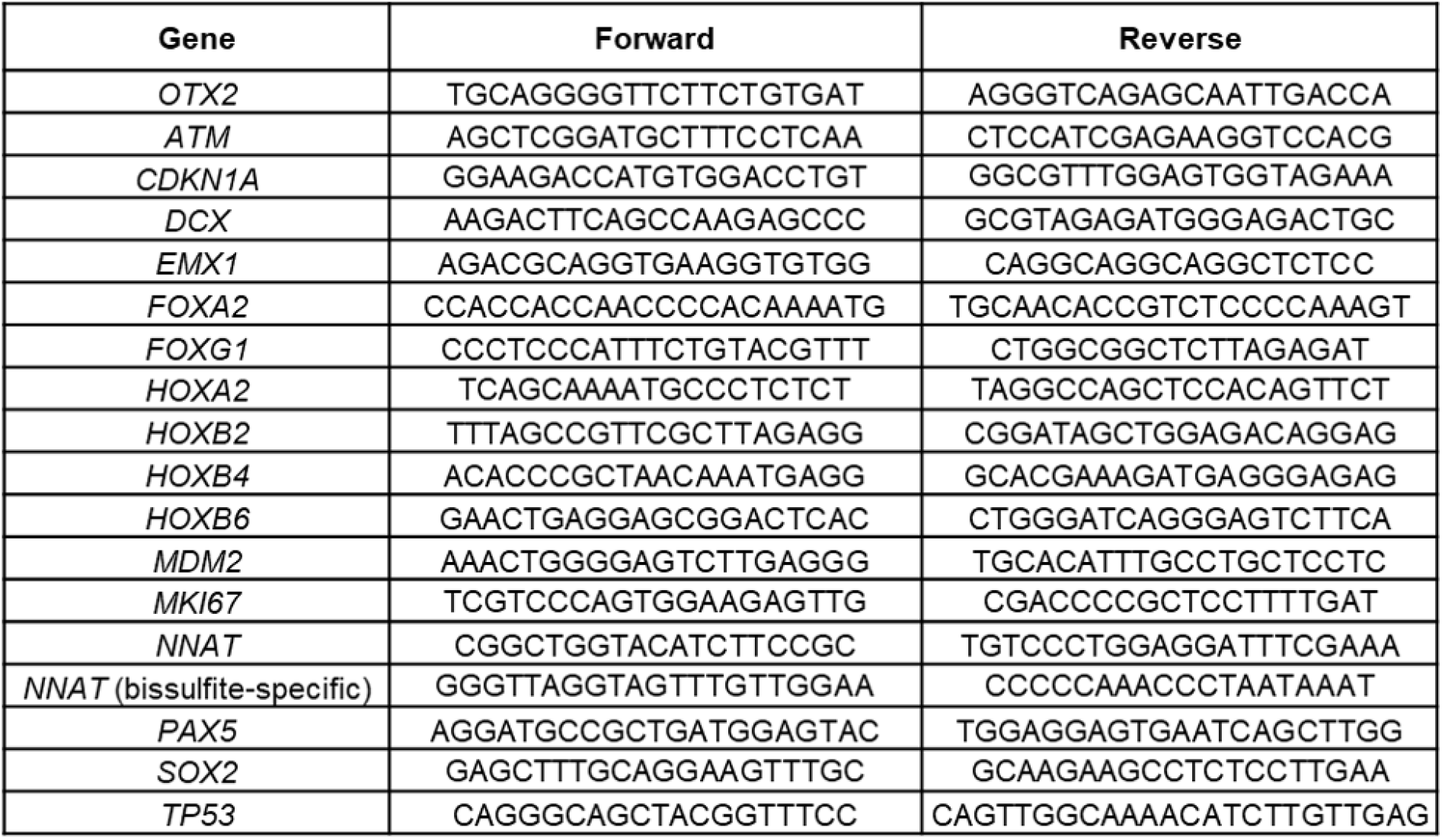
Primer sequences used in this study.

### Supplementary Figures

**Figure S1.**
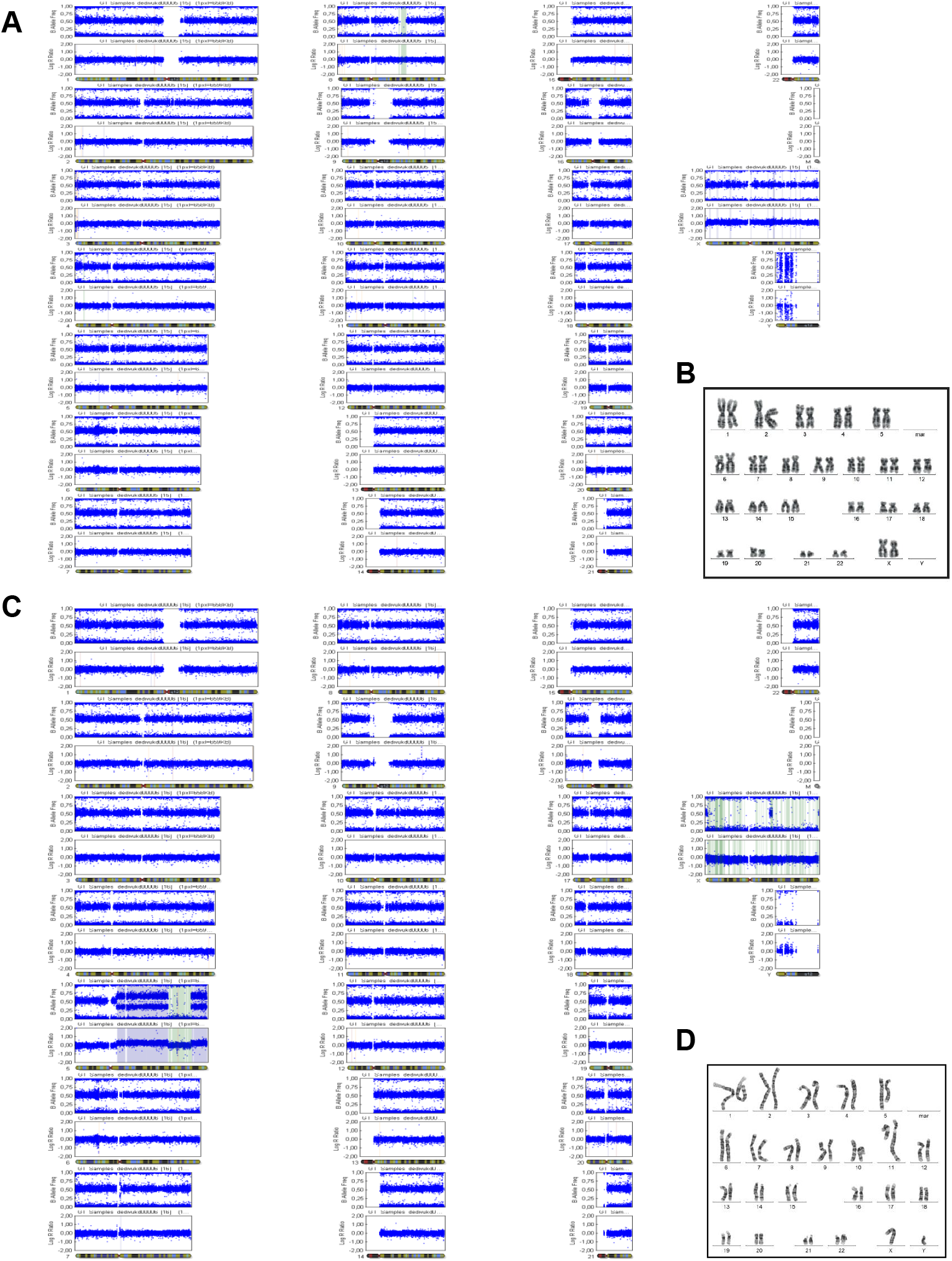
Identification of chromosomal aberrations in NBS-iPSCs. Relative to Table S1. (A) Illustration of whole genome profile of CNV analysis after Array-CGH of NBS1-iPSCs showing a LOH in chromosome 8q. CNV, copy number variations; LOH, lost of heterozygosity. (B) G-banding karyotype of NBS1-iPSC: 46,XX (C) Illustration of whole genome profile of CNV analysis after Array-CGH of NBS8-iPSCs. CNV, copy number variations (D) G-banding karyotype of NBS8-iPSCs: 46,XY,add(5)(q22),der(11)t(5;11)(q14;q?23)[14]/46,idem,add(4)(q31)[10].

**Figure S2.**
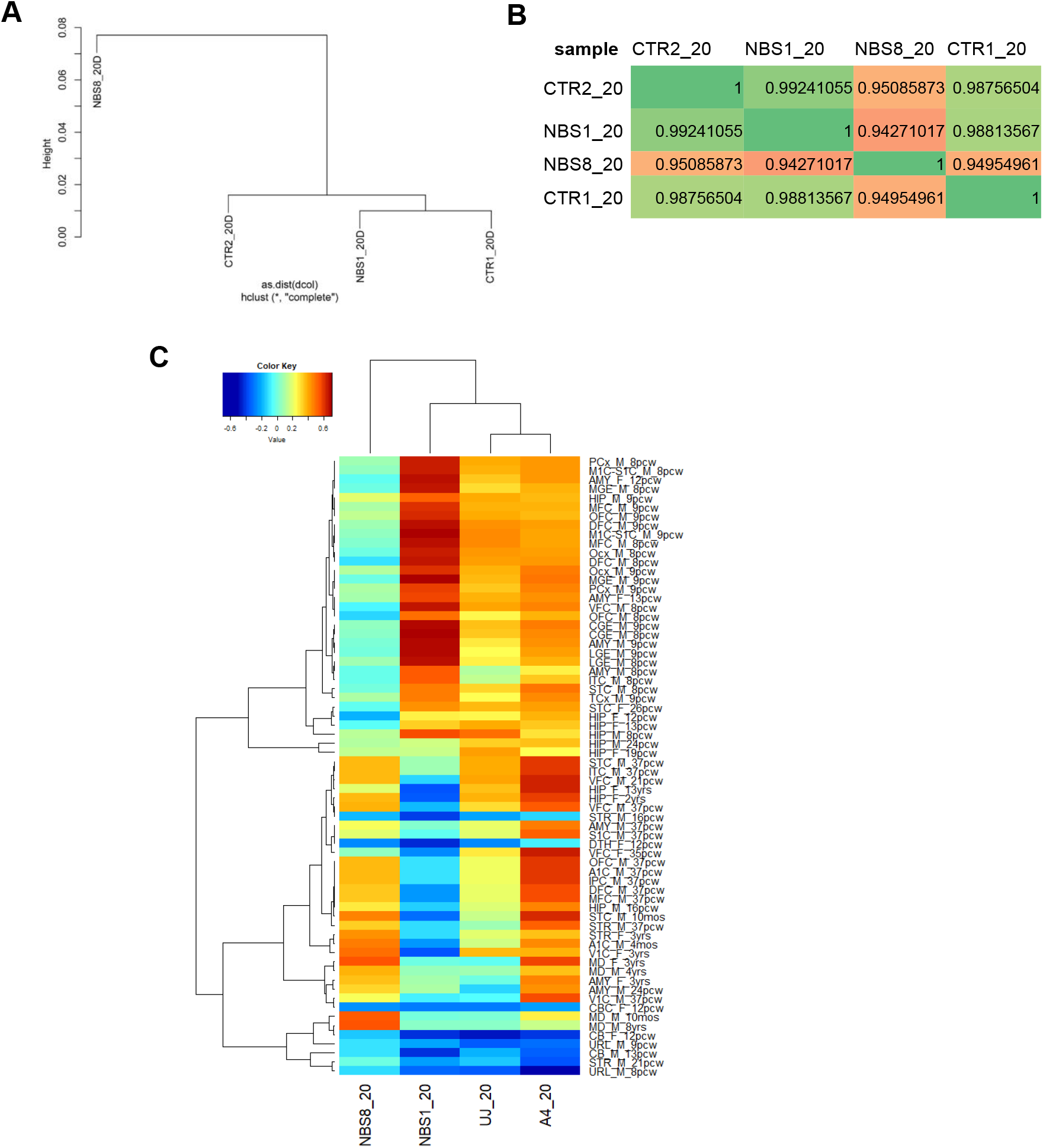
Gene expression profiles of control and NBS organoids. Relative to Figure 3 and Figure 5. (A) Dendogram obtained by hierarchical cluster analysis of microarray-based gene expression data for CTR1, CTR2, NBS1 and NBS8 cerebral organoids. (B) Pearson‘s correlation coefficient matrix of the transcriptomic data showing NBS8 has a distinct transcriptome profile. (C) Heat map of enrichment scores of transcriptomic data from control and NBS organoids in comparison with the Allen Brain Atlas (ABA) human developmental Brain NGS data (https://www.brainspan.org/).

**Figure S3.**
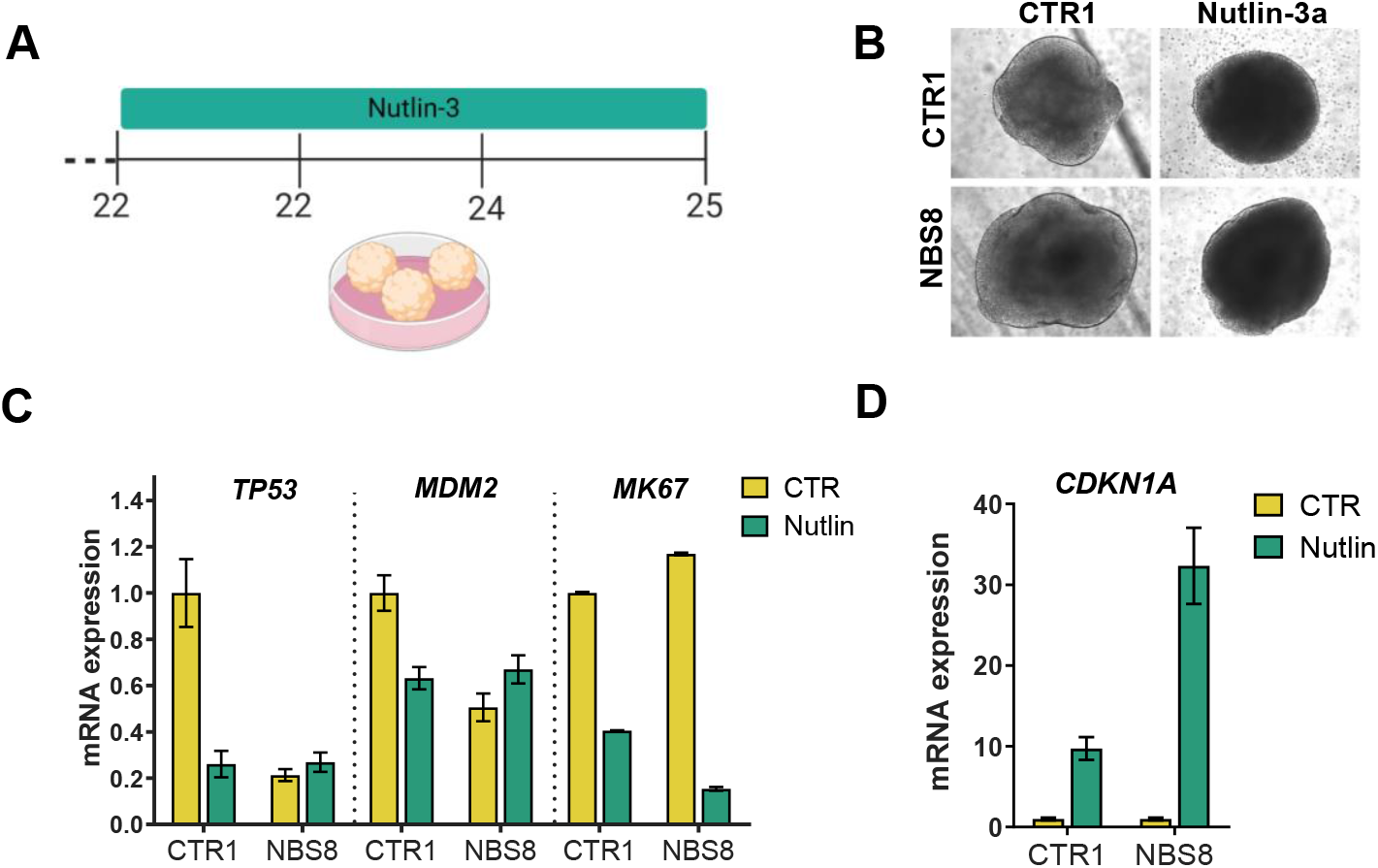
Stabilization of p53-induced by Nutlin 3a. Relative to Figure 4. (A) Schematic depicting the strategy to stabilize p53 via incubation with Nutlin-3a during 72h from day 22 to day 25 of CTR1- and NBS8-organoids. (C-D) qRT-PCR analysis of TP53, MDM2, MKI67 (C) and CDKN1A (D) mRNA expression in CTR1- and NBS8-organoids after Nutlin-3a treatment relative to control (untreated) CTR1: n=3 and NBS8: n=3 technical replicates. Results are mean +/− 95% confidence interval.

**Figure S4.**
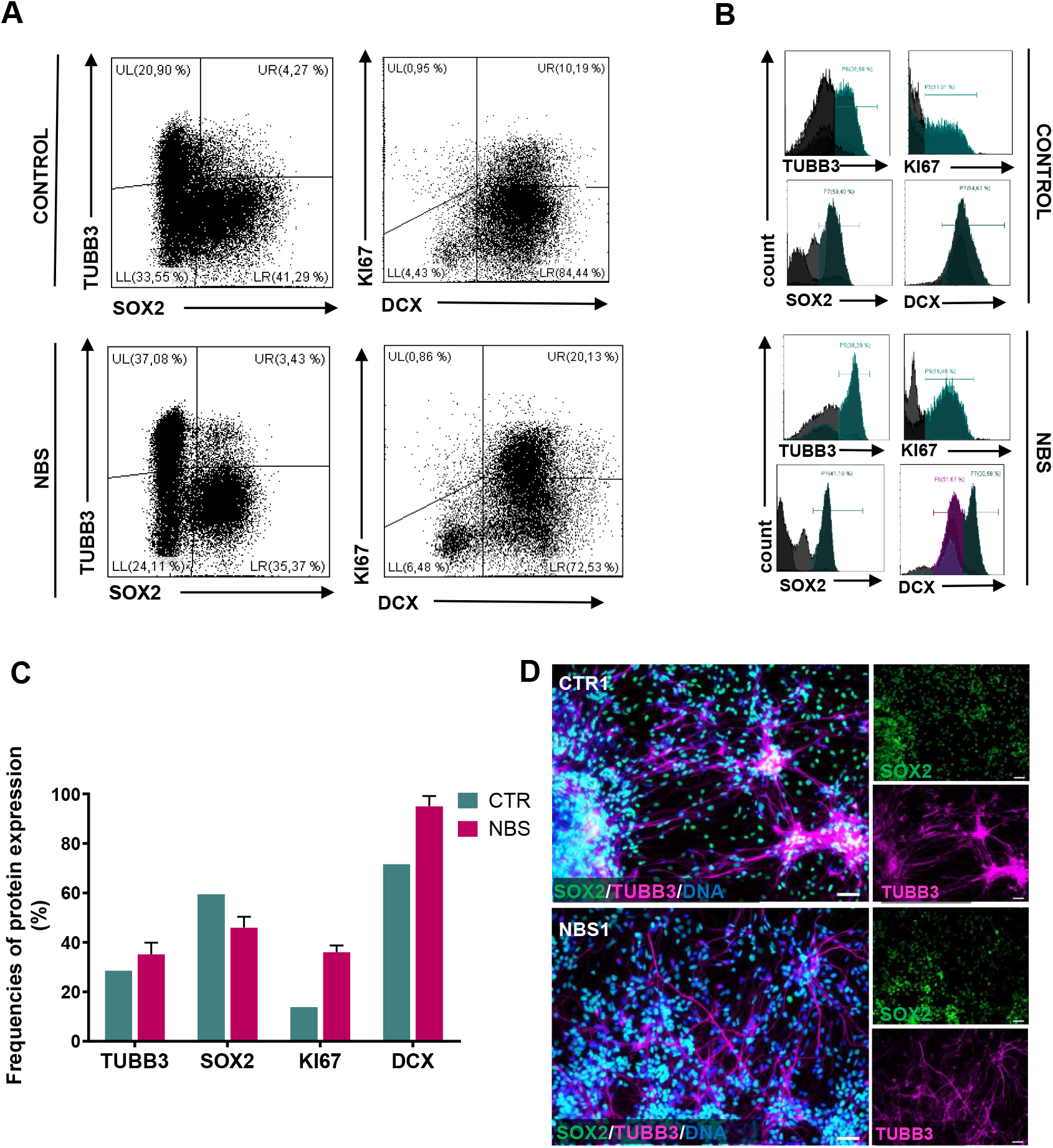
Increase abundance of BIII-Tubulin and DCX-positive neurons in NBS-organoids. Relative to Figure 6. (A) Expression of BIII-Tubulin (TUBB3), SOX2, KI67 and DCX was assessed in single cell suspension of brain organoids by multicolor flow cytometry. Gates and quadrants were set according to negative controls and frequencies of single- and double-expressors are indicated in these representative dot plots. (B) Representative histograms of single-expression of TUBB3, SOX2, KI67 and DCX in single cell suspension of brain organoids. Gates were set according to negative controls. (C) Quantitative assessment of the protein expression in frequencies (%) Data are representative of n 3 independent experiments and presented as mean SD. (D) Representative pictures of immunostainings of SOX2 and βIII-Tubulin in CTR1- and NBS1-organoids after 1 week of single cell dissociation and re-plating in 2D.

**Figure S5.**
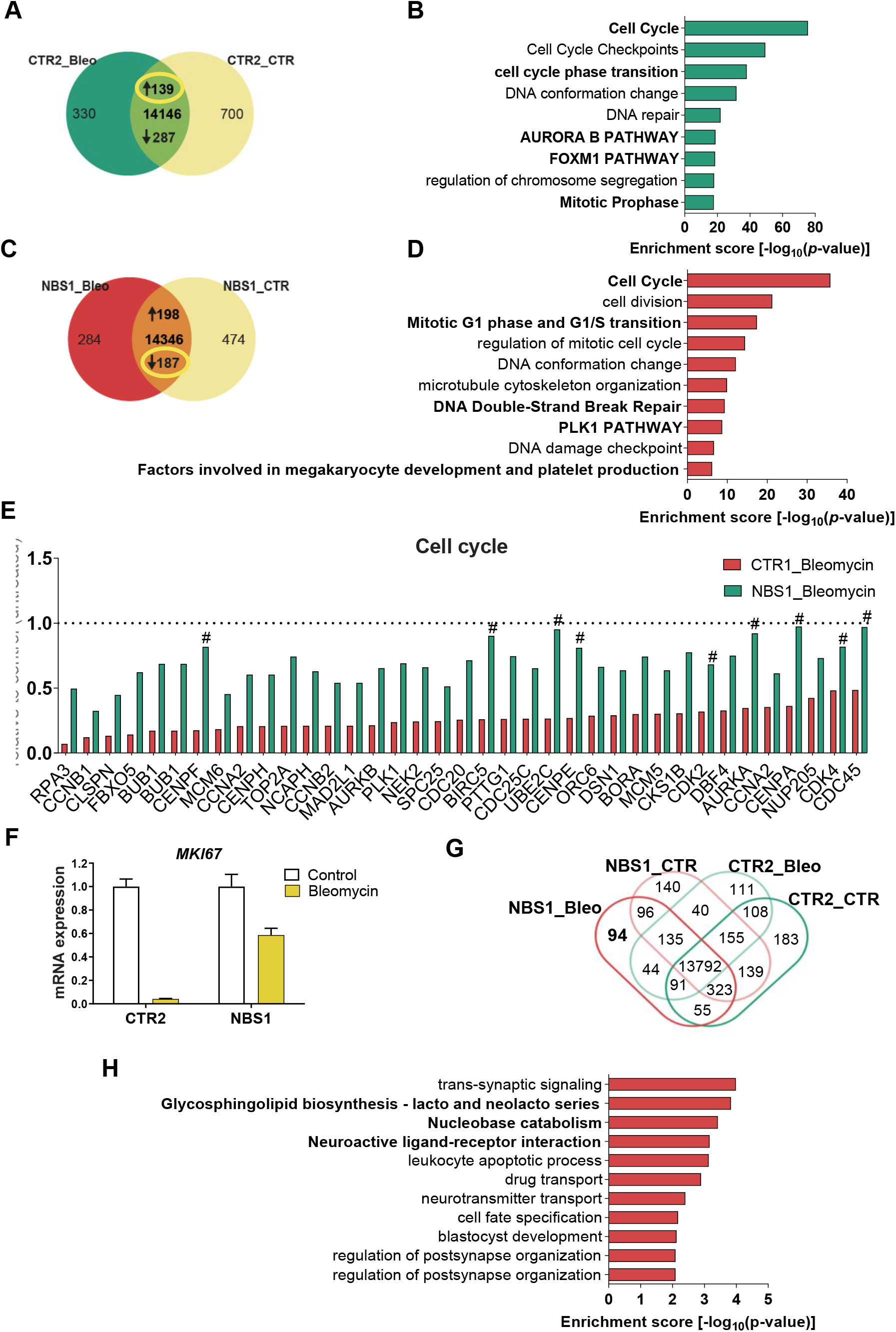
Analysis of the effect of bleomycin in the down-regulated and exclusively expressed genes in CTR2- and NBS1-organoids. Relative to Figure 7. (A-B) Venn diagram showing the 287 down-regulated genes (A) in CTR2_Bleomycin-organoids compared to CTR2_control-organoids and the respective bar chart (B) of the enriched clustered GOs and pathways (Top 10 ranked). (C-D) Venn diagram showing the 187 down-regulated genes (A) in NBS1_Bleomycin-organoids compared to NBS1_control-organoids and the respective bar chart (B) of the enriched clustered GOs and pathways (Top 10 ranked). mRNA expression of genes associated with the down-regulated cell cycle pathway in CTR2_Bleomycin-organoids and NBS1_Bleomycin-organoids compared to CTR_control-organoids and NBS1_control-organoids, respectively. All genes were significantly down-regulated in NBS1 and CTR2 after bleomycin treatment, except genes assigned with in CTR2_Bleomycin-organoids. (F) qRT-PCR analysis of MKI67 of mRNA expression in CTR1- and NBS8-organoids after bleomzcin treatment relative to control (untreated) organoids. CTR2: n=3 and NBS1: n=3 technical replicates. Results are mean +/− 95% confidence interval. (G) Venn diagram dissecting 94 exclusively expressed genes expressed in NBS1_Bleomycin-organoids. (H) Bar chart of the enriched clustered GOs and Pathways (Top 10 ranked) 94 exclusively expressed genes in NBS1_Bleomycin-organoids.

## Notes

### Competing Interest Statement

The authors have declared no competing interest.

